# Digital twin predicting diet response before and after long-term fasting

**DOI:** 10.1101/2021.11.04.467307

**Authors:** Oscar Silfvergren, Christian Simonsson, Mattias Ekstedt, Peter Lundberg, Peter Gennemark, Gunnar Cedersund

## Abstract

Today, there is great interest in diets proposing new combinations of macronutrient compositions and fasting schedules. Unfortunately, there is little consensus regarding the impact of these different diets, since available studies measure different sets of variables in different populations, thus only providing partial, non-connected insights. We lack an approach for integrating all such partial insights into a useful and interconnected big picture. Herein, we present such an integrating tool. The tool uses a novel mathematical model that describes mechanisms regulating diet-response and fasting metabolic fluxes, both for organ-organ crosstalk, and inside the liver. The tool can mechanistically explain and integrate data from several clinical studies, and correctly predict new independent data, including data from a new clinical study. Using this model, we can predict non-measured variables, e.g. hepatic glycogen and gluconeogenesis, and we can quantify personalized expected differences in outcome for any diet. This constitutes a new digital twin technology.

## Introduction

Metabolic dysregulation and obesity lead to many of our most serious diseases: type 2 diabetes, non-alcoholic steatohepatitis (NASH), and cardiovascular diseases. Prevention of these diseases involves proper diet and exercise. Proper dieting involves two central steps: i) figuring out which diet scheme is the best for each person, and ii) making people follow these schemes. Unfortunately, we have still not succeeded with neither of these two steps. There is no consensus regarding which of the many proposed diet and fasting schemes is optimal for each person: some restrict fat while allowing for high carbohydrates (HCLF); some do the opposite, i.e. restrict carbohydrates and allow fat (LCHF); some advocate fasting two days a week (5:2); some 16 hours each day (intermediate fasting, IF); and some suggest small frequent meals (SFM) (Ajala et al., 2013, Ge et al., 2020, Ganesan et al., 2018, Chawla et al., 2020, Dashti and Mogensen, 2017, Sylvetsky et al., 2017, Noakes and Windt, 2017, Scholtens et al., 2020). The reason for this lack of consensus from long-term studies may partially be due to a variety of confounding factors that are hard to monitor in large-scale long-term studies. Well-designed short-term studies show clearer effects. However, in these short-term studies, different studies measure different sets of variables, in different conditions, and in different populations. Such disparate datasets cannot naturally be integrated using traditional methods for data analysis. Finally, there are also many problems regarding compliance to these diets: patients may not believe that the prescribed diet works for them, and/or have problems with motivation and long-term adherence (Jaworski et al., 2018, Gibson and Sainsbury, 2017, García-Pérez et al., 2013).

Lately, a new technology has been advocated that potentially can solve both these problems: physiologically based *digital twins (Fig 1A)* (Shamanna et al., 2020, Corral-Acero et al., 2020, Golse et al., 2021, Schwartz et al., 2020). A physiologically based digital twin is a personalized computer model that describes the underlying physiology in a specific person or patient (Fig 1B). The idea is that you can give your digital twin a meal and fasting schedule, and see how it responds; both acutely to each meal, and etiologically over longer periods of time. The physiological nature of these twins implies that they can deal with the first problem mentioned above: lack of consensus. Specific aspects from each study can be incorporated into the appropriate parts of the twin models (Fig 1C) (Nyman et al., 2016). For instance, a study that measures the interplay between glucose, insulin, and glycogen, will inform and update those aspects of the twin. Another study that measures other variables will update other aspects of the same twin technology. Finally, a digital twin technology can potentially also help with the second problem: lack of compliance. By seeing your own digital twin improve, maintain, or lose its metabolic function by following different diets, the motivation to follow the right diet will likely increase (Fig 1A) (Shamanna et al., 2020).

**Figure 1.**
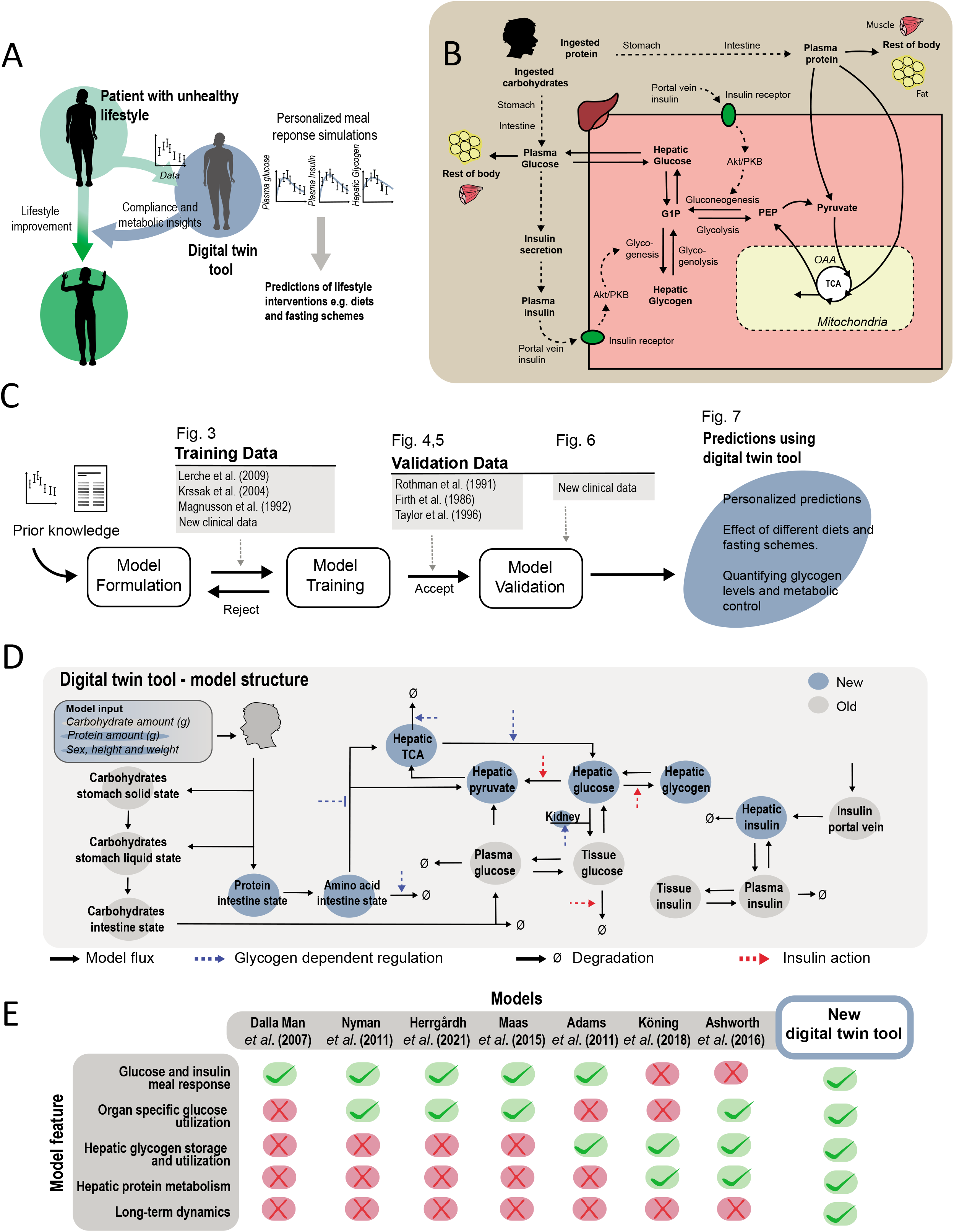
Overview of Digital Twin tool. (A) Illustration of the idea with a digital twin: use personalized predictions and visualizations to increase the metabolic insights and compliance of a patient to prescribed life-style changes. (B) The physiological and biochemical processes considered herein. (C) Workflow and steps taken in this work. (D) Overview of all reactions and regulations included in the model. (E) Comparison between our new digital twin tool and previous physiologically based meal simulation models

There are some well-established physiological models in use for specific types of clinical applications, but no available model can deal with changes between different diet compositions and fasting schemes. While mathematical modelling of meal responses started already in the 1970s (Bergman et al., 1981, Cobelli et al., 1986, Cobelli and Thomaseth, 1987, Bergman et al., 1979), a big breakthrough came with the so-called Dalla Man model in 2007 (Dalla Man et al., 2007). This model is based on data from a triple-tracer experiment, where different glucose and insulin fluxes in the body have been measured. A version of this model is approved by the Food and Drug Administration (FDA) in the US, as a replacement for animal experiments when certifying insulin pumps used to treat type 1 diabetes (Kovatchev et al., 2009). However, while there have been much work improving such models, there is still no model available that: i) can describe the physiological response to different diet compositions containing e.g. different amounts of proteins and carbohydrates, or ii) can correctly predict the response to fasting (Fig 1E). For these reasons, the potential of a digital twin mentioned above cannot yet be realized. In this paper, we resolve both of these shortcomings. Using four different clinical studies (Fig 1C, training data, Fig 2–3), we develop and train a new and significantly extended model (Fig 1D). The model can predict new personalized responses (Fig 1C, validation data, Fig 4–5) from both previous studies, and from a new study involving oral protein tolerance tests (OPTT) before and after 48 h fasting (Fig 6). Finally, we illustrate that the combined knowledge extracted from all of these studies can be used to predict the patient-specific impact of a variety of dietary compositions and fasting schedules (Fig 7).

**Figure 2.**
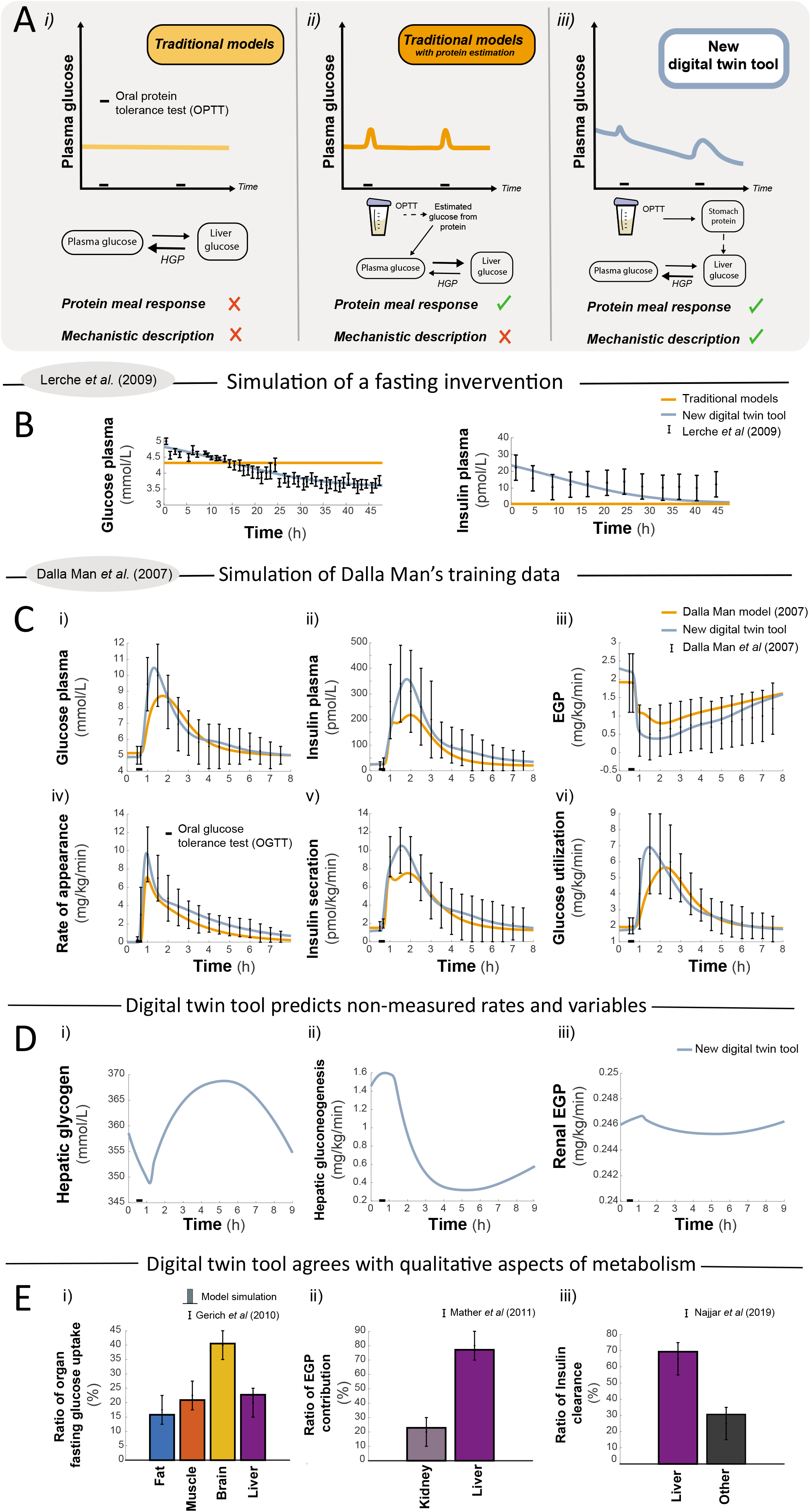
Model features compared to previous model. (A) Overview of the qualitative predictions and improvements in our model, compared to previous models. Previous models can only either i) ignore protein ingestion in glucose simulations, or ii) have a static formula for a phenomenological conversion between protein and plasma glucose, which incorrectly increases the hepatic glucose uptake. In contrast, iii) in our new model, we can have a protein-induced increase in gluconeogenesis and hepatic glucose production (HGP), which is much larger after fasting than in a fed state. The arrows and their sizes represents the qualitative relationships between the fluxes between plasma glucose and plasma liver during the second OPTT response, for the three model alternatives. (B) Experimental data (error bars) validating another key difference between the new (blue line) and old model simulations (orange line): the decrease of plasma glucose during fasting. (C) Our model (blue line) can describe all of the mechanistic flux data (error bars) that led to the Dalla Man model (Dalla Man et al., 2007), equally well as that model and its subsequent improvements (orange line). Data shows responses to a mixed meal at 0.5 h (black bar)(details of all clinical studies and data in Fig S4). (D) Examples of predictions of key mechanistic variables that our model can produce, which the original Dalla Man model and its subsequent improvements (including (Herrgårdh et al., 2021)) cannot produce. (E) Our new model also produces organ-specific predictions (filled bars) which agrees well with data (error bars) for glucose uptake (i, (Gerich, 2010)), endogenous glucose production (EGP) from liver and kidney following a meal (ii,(Mather and Pollock, 2011)), and insulin clearance (iii, (Najjar and Perdomo, 2019)).

**Figure 3.**
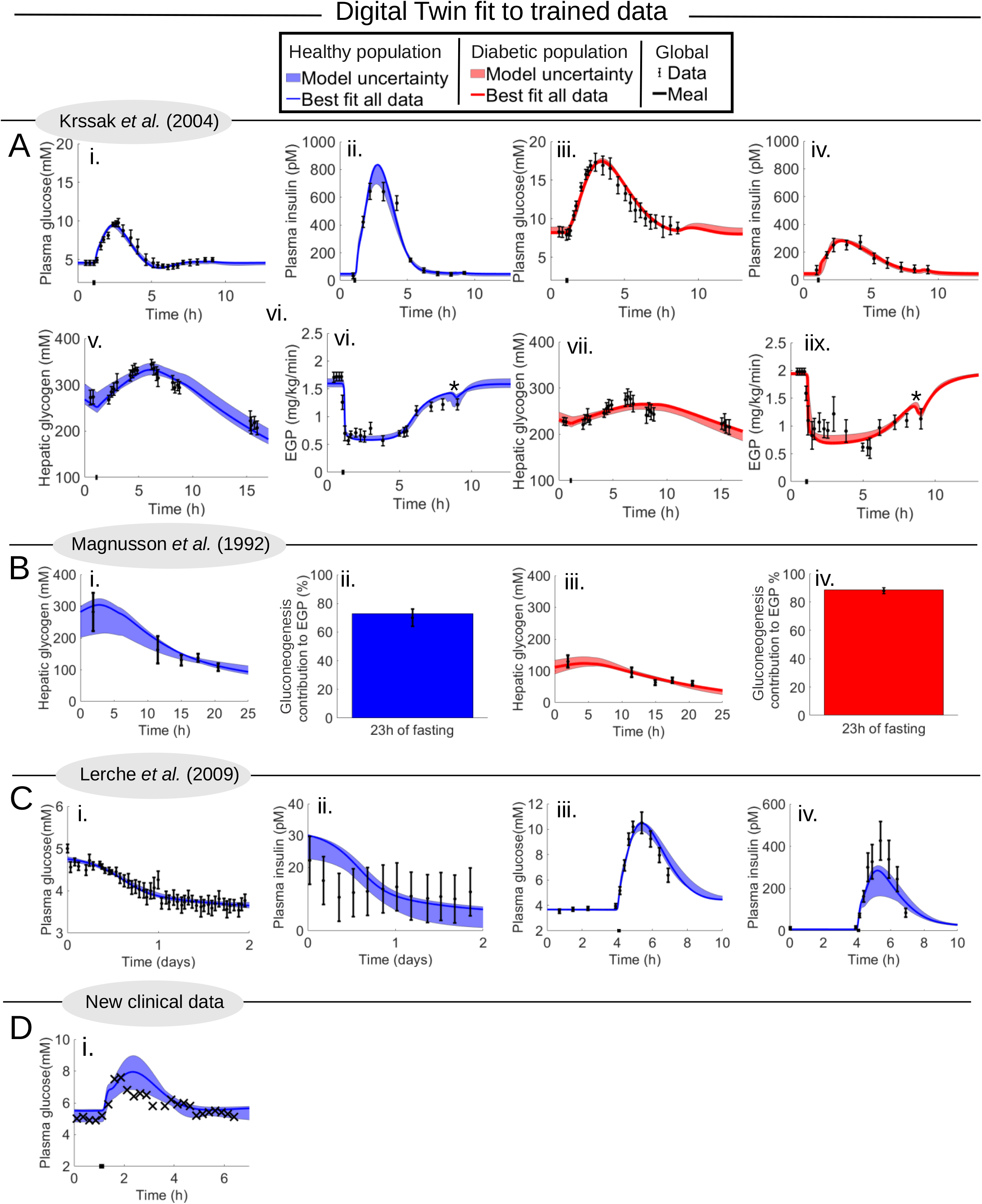
Model simulations of the four clinical studies used for parameter estimations. In all plots, datapoints are represented as the centre of error bars (SEM) or crosses. Simulations with uncertainty are represented by areas, and the best agreeing simulation is the line, for both healthy populations (blue) and T2D population (red). Quantitative details regarding all studies are summarized in Table S4. All of these data are mean responses, and simulations are made for a 175 cm, 75 kg male person. (A) Krssak data (Krssak et al., 2004), which describes a mixed meal response (87 g carbohydrates and 23 g protein) happening at 1 h (black bar), whereafter plasma glucose, insulin, and EGP and hepatic glycogen are measured. (B) Magnusson data (Magnusson et al., 1992), which describes a fasting response, following a mixed meal (98.2 g carbohydrates and 26 g protein) which happens 4 h before t=0. (C) Lerche data (Lerche et al., 2009), which describes a 48 h fast, which starts at t=0, which is preceded by an overnight fast, and which is followed by an OGTT of 1 g carbohydrate/total bodyweight [kg]. In iii) and iv), the time has been shifted, so that the meal occurs at t=3.5 h. (D) Our new clinical study, which in this training data describes the glucose response to a mixed meal (81 g carbohydrates and 41 g protein, in total 940 kcal) at t=1 h (black bar).

**Figure 4.**
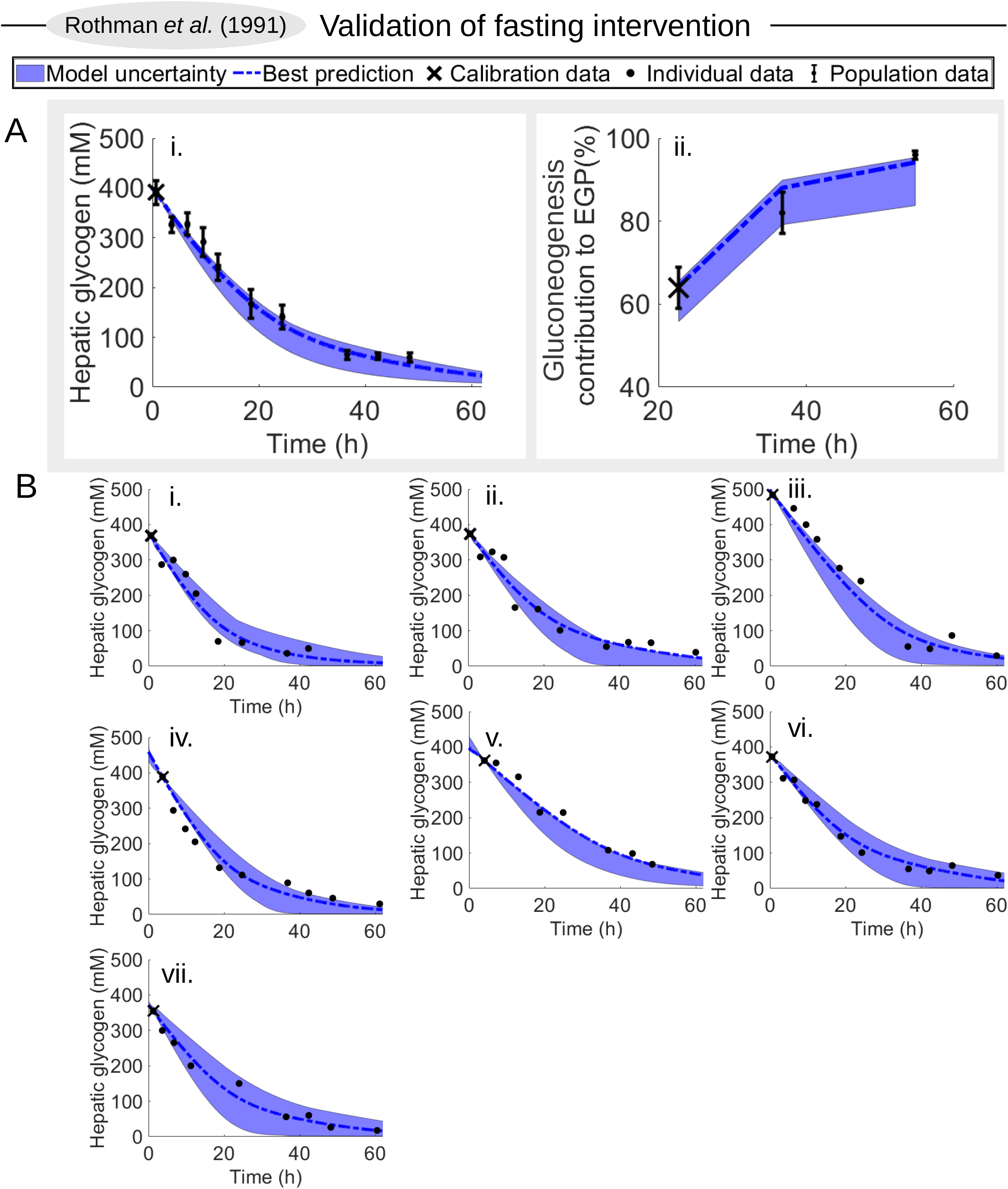
The first validation test done with the model, using the Rothman data (Rothman Douglas et al., 1991). In all plots, data values are depicted with a dot, SEM uncertainty by the error bar, simulation uncertainties are depicted by the areas, and a simulation using the model parameters that best agree with the data are depicted with the line. The first datapoint (also depicted with an X) is used for personalizing the model (Supplementary material), and the rest are used for validation. The study monitors the response of hepatic glycogen, and of the contribution of glycogenolysis to EGP, during a 68 h fast. (A) Prediction of hepatic glycogen levels and gluconeogenesis contribution to the endogenous glucose production on the full population. (B) Prediction of hepatic glycogen levels on an individual level.

**Figure 5.**
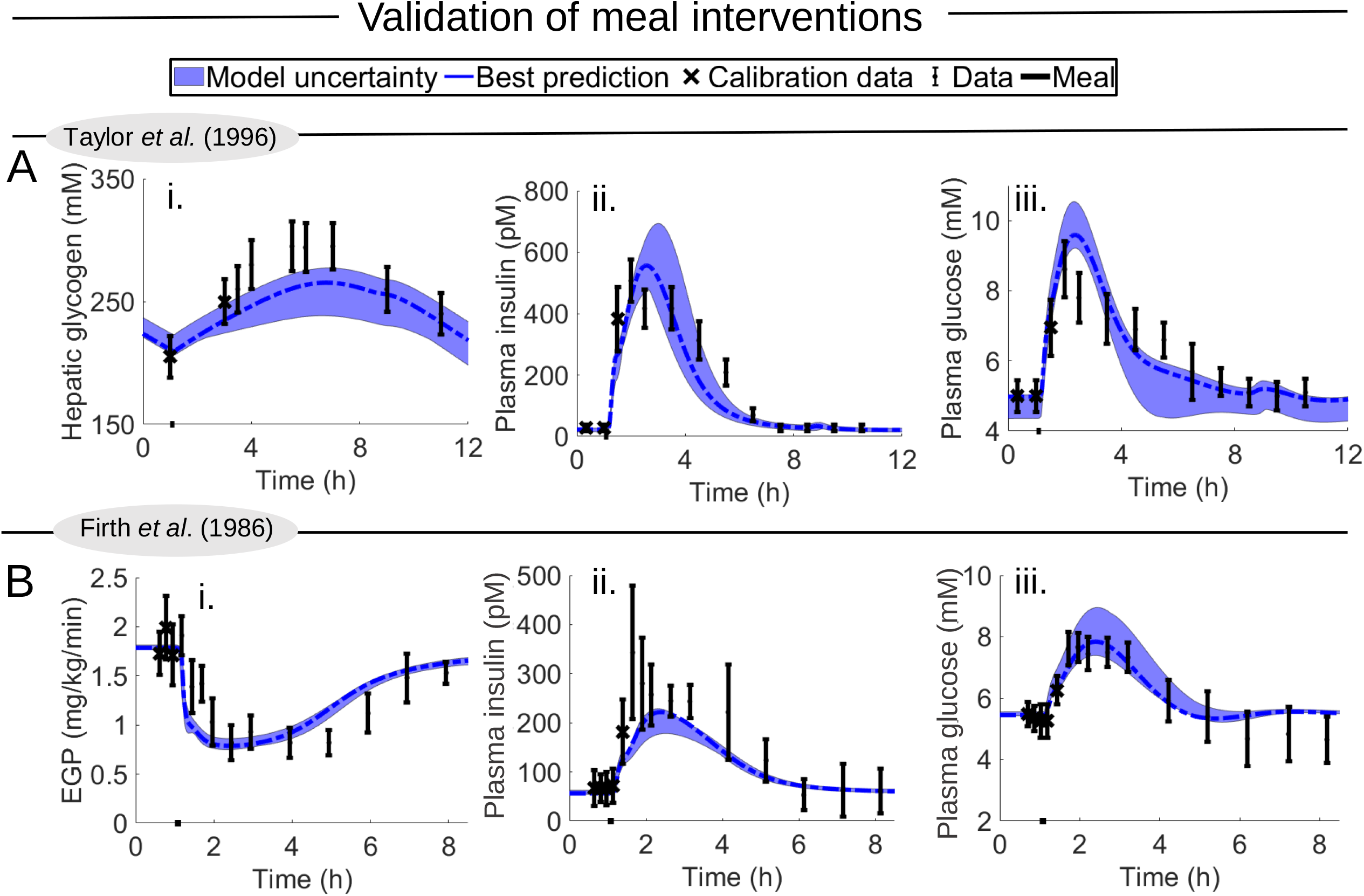
Second validation test with the model, using two clinical studies not used for training the model (Taylor et al., 1996, Firth et al., 1986). In all plots, data values are depicted with error bars (SEM), simulation uncertainties are depicted by the areas, and the simulation that best agrees with the data is depicted with the dash-dotted line. The first couple of datapoints (depicted with an X) are used for personalizing the model (Supplementary material), and the remaining data points are used for validation. (A) Prediction of mixed meal (black bar) from Taylor data (32), i) hepatic glycogen ii) plasma insulin iii) plasma glucose (B) Prediction of OGTT (black bar) from Firth data (32), i) endogenous glucose production ii) plasma insulin iii) plasma glucose Additional details regarding each study are summarized in Table S4.

**Figure 6.**
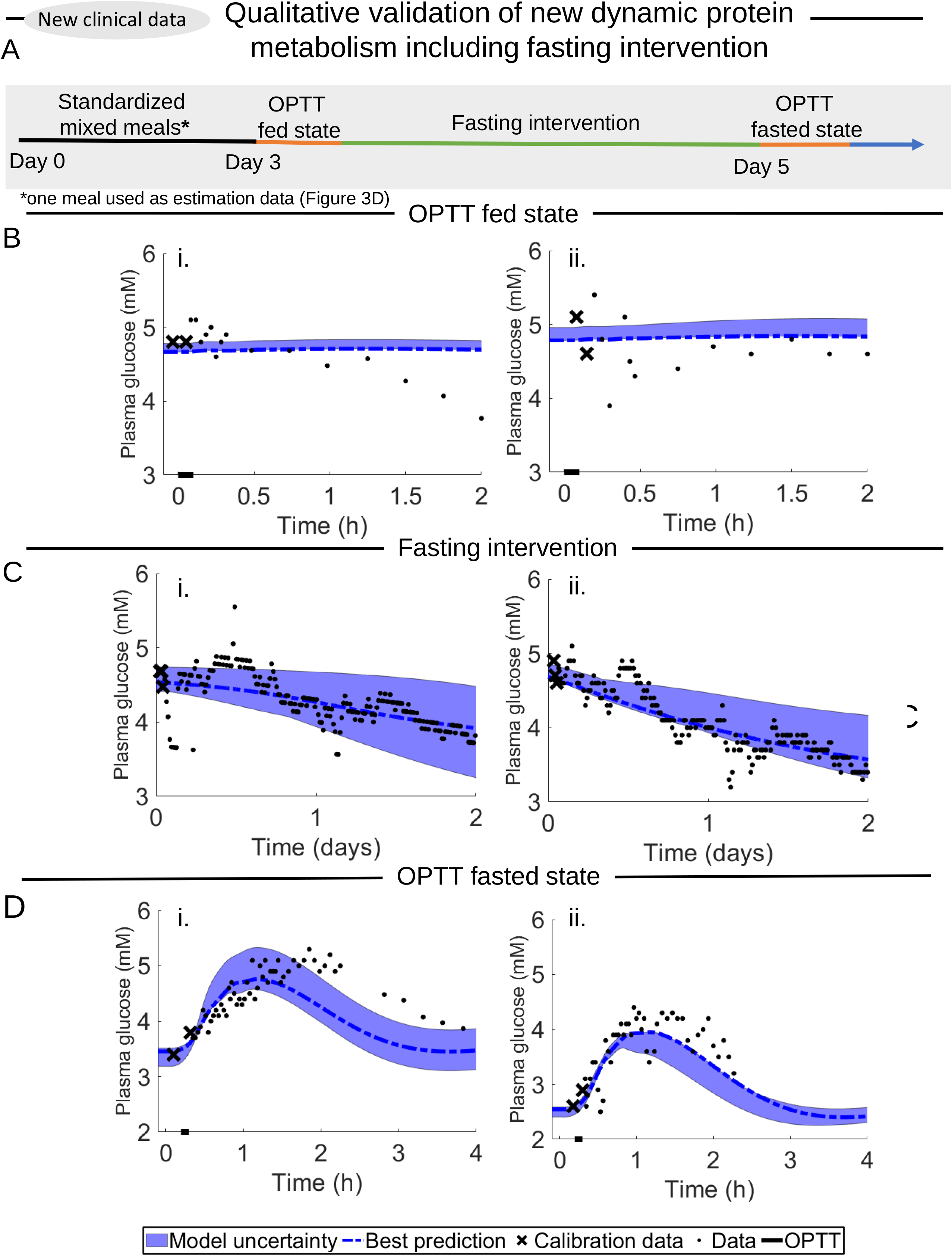
third validation test of new data testing model’s ability in predicting fasting intervention and metabolism of proteins. Model uncertainty is depicted with the area, the calibration data is depicted with an X, the calibrated CGM data are depicted with a dot, and time and duration of the OPTT is depicted with a black line. (A) Study design. (B) OPTT response during the fed state, before the fast started. (C) Glucose levels during the 48 h fast. (D) OPTT response during the fasted state, after the 48 h fast.

**Figure 7.**
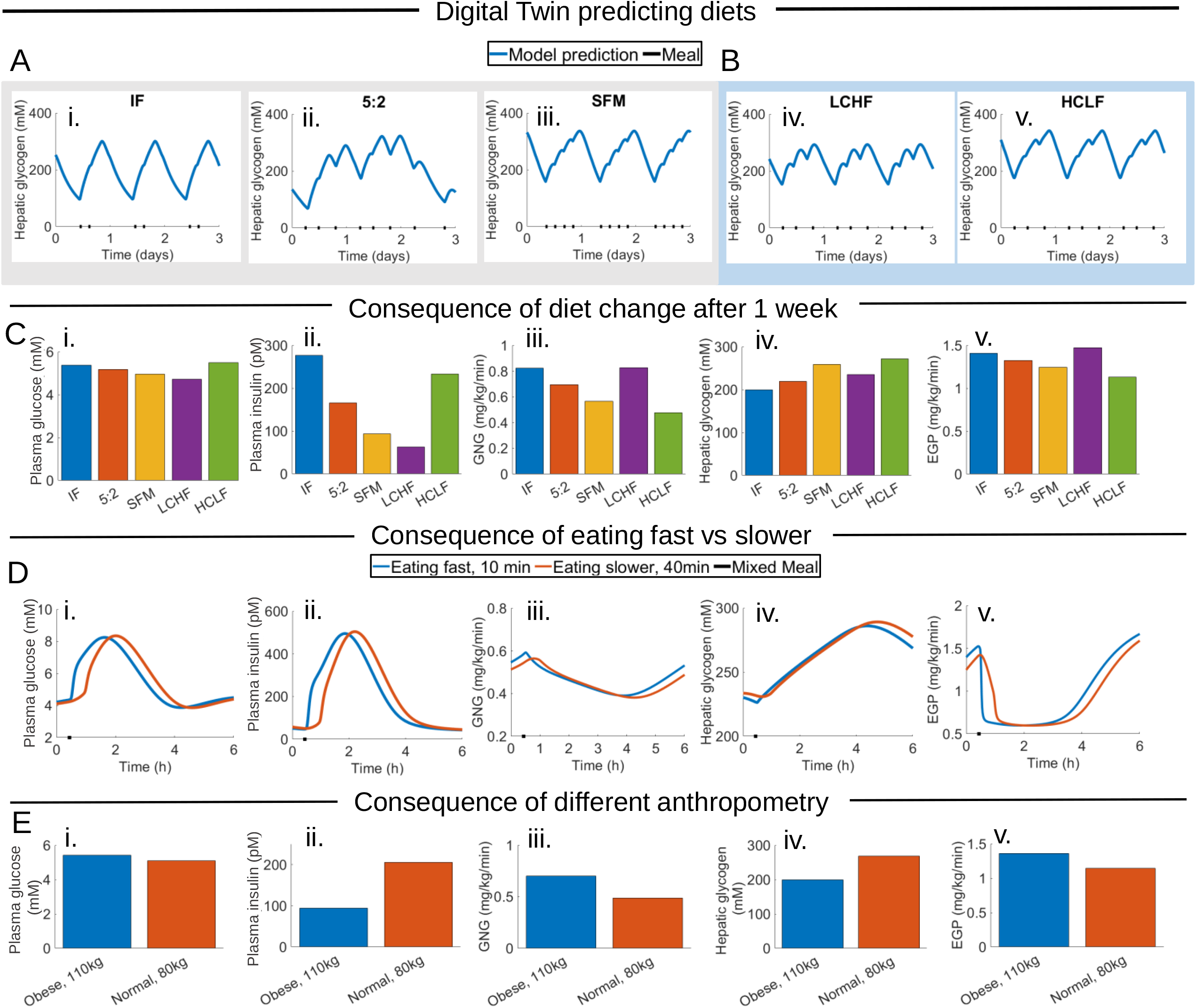
simulations 5 common diet schemes and varying different personalized variables. (A) The model considers diets with different meal frequencies, ranging from Intermediate Fasting (IF) with two meals (depicted as black bars), to 5:2 with 2-3 meals, and even SFM (with three smaller meals). These three diets are isocaloric. (B) The model can also to some extent simulate different compositions, such as LCHF (low carbohydrate, high fat diet), and HCLF (high carbohydrate, low fat diet). (C) Mean values of key variables during the second week after the diet has started. (D) Impact of changing the rate of consumption. E) Impact of changing the body weight by 30 kg, while keeping all other variables constant. All simulations in A-D are done using a male who is 180 cm, 80 kg, and in E that male either weighs 110 or 80 kg.

## Results

### We have substantially improved and expanded the capabilities of the previous meal models

We have extended our most recent improved version of the original Dalla Man model (Herrgårdh et al., 2021), to include a series of new features (Fig 1D, blue): i) intracellular metabolism in the liver, ii) long-term energy regulation via the new states for liver and kidney glycogen, iii) protein metabolism, and iv) hepatic interconversion between glucose and amino acids. Because of these changes, our digital twin tool can incorporate data and insights from a wide variety of studies, measuring e.g. fasting, glycogen, gluconeogenesis, and both hepatic and kidney endogenous glucose production. Our model has been iteratively developed and modified (Fig 1C) to be able to describe the variety of studies used in the estimation data (Fig 3). The resulting model is fully described in detail in the 25+ page long Supplementary material, and all model files are available on github (Material and methods). The reliability of our new tool is tested using new data (Fig 4–6). Both estimation and validation data include various aspects of a new clinical study, which has been carefully designed to generate new responses not present in any of the other data: responses to large meals, in the estimation data (Fig 3D), and oral protein tolerance test responses before and after 48h fasting, in the validation data (Fig 6). Finally, because of these new improvements, the model can simulate physiologically based predictions of diet-responses (Fig 7), which both constitutes an additional qualitative validation, and which illustrates how the model can be used in clinical applications: to serve as a digital twin that provides personalized diet simulations and thus may be used as a tool in helping with metabolic dysregulation and obesity.

### Digital twin tool can describe all original data used to develop the Dalla Man model

First, to check that our model has not lost any critical ability compared to its predecessors, the new model was fitted to the original data used to develop the Dalla Man model (Dalla Man et al., 2007) (Fig 2C). In that study, healthy subjects were subjected to a meal response at t = 0.5 h (black line), and a triple-tracer protocol was used to measure a variety of variables: plasma glucose, plasma insulin, endogenous glucose production (EGP), glucose rate of appearance, glucose utilization, and insulin secretion. Compared to the previous model (orange line), our new simulations (blue line) agree at least as well with the data (error bars). In fact, for some variables, such as plasma insulin and glucose clearance, the agreement with data is enhanced using our new model. However, the exact parameters used in the original model publication are not publicly available, and the orange line depicts simulations with parameter values we have used in previous publications, where we analyzed the Dalla Man model (Sips et al., 2015, Herrgårdh et al., 2021).

A more important improvement in these curves concerns the underlying mechanisms for some of the simulated dynamics, e.g. EGP. In the original Dalla Man model, the drop in EGP following a meal is explained by a phenomenological expression, which simply subtracts the rate of appearance flux from a constant value. Our model simulations produce the same drop in EGP, but using mechanistic processes, involving e.g. insulin inhibition of gluconeogenesis and glycogenolysis (Fig 1D).

Finally, that the agreement between data and simulation is acceptable is supported by both visual assessment and by a *χ*^2^-test 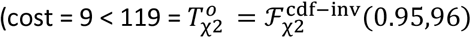, Materials and methods).

### Our digital twin tool can describe data for short and intermediate-term dynamics in glucose, insulin, EGP, glycogen, and gluconeogenesis, both in healthy and T2D populations

The model was trained on data from four different clinical studies (Fig 3), three already published (Lerche et al., 2009, Krssak et al., 2004, Magnusson et al., 1992)(Fig 3A–C) and a meal from our new study (Fig 3D). These studies include various measured variables in both healthy and diabetic conditions, and the model can successfully describe all these studies. The model is trained to all healthy data together, and to all diabetic data together, but all parameters are allowed to be different between healthy and diabetic conditions. Apart from this, each study considered in this paper is modified according to some covariates: age, weight, height, and sex (Supplementary material S4). The agreement between data (error bars) and simulations (areas) is acceptable according to both visual assessment and a *χ*^2^-test, for both healthy data 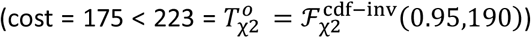 and for diabetes data 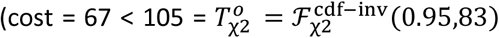, Materials and methods). This means that the model provides an acceptable mechanistic explanation for all of these data (Cedersund and Roll, 2009).

The first study (Krssak et al., 2004)(Fig 3A) shows mixed-meal responses for hepatic glycogen, glucose, EGP, and insulin, in a healthy (blue) and a diabetic (red) population. The meal is given at t = 1 h, represented with a black bar along the x-axis. Glucose and insulin reach a peak around 1 h, and returns back to a baseline within 5 h. In contrast, the dynamics of glycogen is slower, with a peak around 2.5 h, and the concentration is still declining after 9 h. As can be seen, the model simulations (areas) replicate these qualitative features, and in general lie close to experimental data (error bars) for all variables, which also is supported by the *χ*^2^-test.

The second study (Magnusson et al., 1992)(Fig 3B) looks at two things - the role of gluconeogenesis and the amount of hepatic glycogen - in both healthy and diabetic populations. The subjects eat a standardized meal 4 h before the first sampling point, whereafter they are fasting for 23 h (Table S4). The authors estimated the contribution of gluconeogenesis directly from data by subtracting measured EGP from decrease in glycogen. In the model, this difference, that is the contribution of gluconeogenesis, corresponds to the sum of gluconeogenesis in both liver and kidneys divided by the total EGP. As can be seen, the contribution of gluconeogenesis is around 70% in healthy individuals, and around 88% in diabetic individuals, in both data and model. This means that diabetic patients rely more on gluconeogenesis, and that healthy individuals rely comparatively more on glycogenolysis. This difference happens because healthy individuals have higher glycogen levels, which are lower in diabetic individuals, because of their insulin resistance and reduced insulin production. All these differences are seen both in the data and in the model.

The third study (Lerche et al., 2009)(Fig 3C) again considers a fasting intervention, this time 48h, following an overnight fast. Both the model (area) and data (error bars) display a drop in glucose during these 48 h (Fig 3C,i). During this time, the insulin drops from around 30 to around 10 pM in the model. While this drop in insulin lies within the experimental uncertainty, the drop as such is not as apparent in the data. This possible discrepancy has to do with the insulin production part of the model, which is partly dependent on the current glucose concentrations in plasma. This part of the model dates back to the original Dalla Man model (Dalla Man et al., 2007) (see Supplementary file for a more extensive explanation of how model works). After this fasting period, an oral glucose tolerance test (OGTT) of 300 kcal is performed. In Fig 3C,iii,iv, this OGTT is ingested at 3.5 h, and gives rise to a glucose peak of around 10 mM, which is higher than normal in a healthy population, because of the prior long fasting period. As can be seen, the model describes this unusually large OGTT response for both glucose (Fig 3C,iii) and insulin (Fig 3C,iv).

The final fourth study, concerns data on a meal response, taken from a new clinical study conducted by us (Fig 3D). The reason for this addition is that all other estimation data has a relatively small meal size (<450 kcal). Since we want the model to be able to describe meal responses during normal life, which often goes up to ~1000 kcal, we included one of the meals from the first five days in our clinical validation study (Fig 6) also in the estimation data. This additional training data describes glucose dynamics in a mixed meal of ~940 kcal. As can be seen, the glucose peak is around 8 mM in both data and model.

In summary, all of these simulation results show that the model has the ability to find a mechanistic explanation for both short-, and long-term dynamics in glucose, insulin, EGP, glycogen, and gluconeogenesis, in both healthy (blue) and T2D populations (red).

### Our digital twin tool can correctly predict subject-specific hepatic glycogen levels and gluconeogenesis contribution to endogenous glucose production

The first validation test done with the trained model was with the Rothman data (Rothman Douglas et al., 1991)(Fig 4). In this study, they observed the dynamics during a 68 h fast in a healthy population. The fast started at t=0. On the population level, they measured hepatic glycogen and gluconeogenesis contribution to EGP (Fig 4A). These variables were estimated in the same way as in the Magnusson study (Fig 3B). However, in the Magnusson study, the fast only lasted 23 h, and here it lasted more than twice as long. Furthermore, in this study, individual hepatic glycogen responses were reported for the 7 individual subjects (Fig 4B). As can be seen, the model (area) agrees well with the experimental data (error bars) for all these observations. Here, the personalization of each digital twin is done in two ways: i) covariates age, weight, height, sex, and diabetes status are set to reported values, ii) the meals during the days leading up to the start of the study are optimized to agree with the initial datapoints, marked with an X (Supplementary material). This visual assessment of the acceptable agreement is formally supported by a *χ*^2^-test 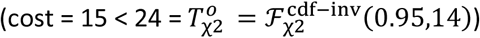.

### Our digital twin tool can correctly predict glucose, insulin, and EGP in independent studies

The second validation test done with the model trained to the data in Fig 3, was done using the data in Fig 5 (Taylor et al., 1996, Firth et al., 1986). This data includes two different studies: one OGTTs (Fig 5A) and one mixed meal (Fig 5B). These two studies were chosen because they measure not only plasma glucose and insulin, but also one additional variable that has to do with the new additions to the model: hepatic glycogen (Fig 5A) and EGP (Fig 5B). Considering the size and meal content are different in both studies, the response in glucose and insulin is qualitatively similar in both studies. EGP declines in response to the OGTT and then slowly goes back to normal, on a timescale that is comparable to the glucose and insulin meal responses. In contrast, the hepatic glycogen is increasing in response to the meal with a peak after about 5-7 h, and hence on a slower timescale than the other variables. All these observations are correctly predicted by the model, both qualitatively and quantitatively. As before, this visual assessment is supported by a *χ*^2^-test, both for the mixed meal 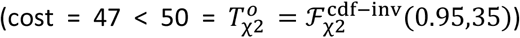 (Fig 5A) and for OGTT 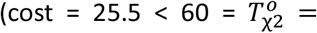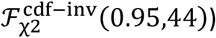 (Fig 5B). Nevertheless, the digital twin tool does predict a slightly lower synthesis of hepatic glycogen from the mixed meal (Fig 5A, i) compared to the experimental data. This slight discrepancy may be a product of underlying differences between the population used in the training of the model and the population used in the Taylor study.

### Our digital twin tool can predict fasting interventions and protein metabolism both in a fed and unfed state

To further test the capabilities of the model, a new dataset was collected, which tests a qualitatively new type of prediction, not present in any of the other datasets. More specifically, the new study examines the response to an OPTT before and after a 48 h fasting (Materials and methods). These combined OPTT and fasting responses are highly central predictions for the new additions to our digital twin model. The reason for this centrality, is that these predictions concern the interplay between glucose and protein metabolism and how this interplay is impacted by glycogen and fasting. These interplays and regulations were not present in previous models (Fig 1E).

Three different predictions were central to the design of the clinical study (Fig 6A). Firstly, plasma glucose will decrease during the 48 h fasting (Fig 2A, iii). This prediction is different compared to the corresponding prediction of the prior meal response models (Fig 2A, i, ii), and it is supported by data (Fig 6C) for both subjects. However, this prediction was identified already in the training data, for example in the Lerche study (Fig 3C). In contrast, the second and third predictions concern the OPTT responses before and after this fasting period, and these predictions have not been seen in any training data. The second prediction is that the first OPTT response, before the fasting, will not give rise to any noticeable response in plasma glucose concentration, and this is what is seen in the data (Fig 6B). In contrast, the third and last prediction is that the same OPTT will give rise to at least 1 mM increase in plasma glucose after the fasting, and that this rise will remain above the glucose values before the OPTT for at least 2 h (Fig 6D). As can be seen, these observations are clearly fulfilled by the data. Since we are now in a digital twin setting, with subject-specific responses, each datapoint comes without an associated uncertainty, and a direct *χ*^2^-test is not possible.

In summary, our new clinical study validates a dynamic and qualitatively novel prediction done by the model.

### Our digital twin tool illustrates qualitatively accurate behaviour of the human metabolism including insulin degradation from liver and plasma, organ specific glucose uptake, and renal glucose production

Unlike the original Dalla Man model, our new model can produce organ-specific predictions (Fig 2E), such as organ-specific glucose uptake; contribution of renal and liver glucose production to EGP; and the ratio of how much insulin is cleared by the liver compared to clearance by the rest of the body. The model simulates reasonable organ-specific glucose uptake both in a post-adsorptive state and postprandial state. In a post-adsorptive state, such as overnight fasting, organ-specific glucose uptake ratios have been measured to be approximately 40-50% in brain, 15-20% in muscle, 10-15% in liver, and 5-10% in kidneys (Gerich, 2010). These data are in good alignment with the model simulations, which lie within these uncertainty regions for all organs (Fig 2E, i). The model can also describe renal glucose production (RGP) in qualitatively accurate proportions compared to hepatic glucose production (HGP). RGP approximately produces 5-10% of all endogenous glucose in a post-adsorptive state and 10-15% in a postprandial state (Gerich, 2010), which is well described by the model (Fig 2E,ii). Finally, the model also simulates a reasonable insulin rate of disappearance produced by the liver, compared to the rest of the body (Fig 2E,iii). Note that these experimental data have a high degree of variation, with values ranging from 80% (Najjar and Perdomo, 2019) to 50% (Duckworth et al., 1998). Note that all organ proportion data referred to come from a variety of different studies, most of which differ to the model simulations considered herein.

### Our Digital twin tool predicts short and long-term responses to common diet schemes

To illustrate the potential of our digital twin tool, we simulate the impact of changing the meal frequencies (Fig 7A), meal compositions (Fig 7B), ingestion speed (Fig 7C), and bodyweight (Fig 7D). The change in diet is performed using a 3-week simulation protocol. The first week is used to reach a steady state using a standardized diet. In the second week, a new diet is introduced, which eventually reaches a new steady state behavior. The third week is the week plotted and analyzed in Fig 7. The plots in Fig A and B depict three days during this third week, to illustrate the daily variations. Note that one of the diets, the 5:2 diet, has different numbers of meals and different meal sizes during the 5 normal days (2560 kcal/day), compared to the 2 restricted days (600 kcal/day). Note also that all diets in Fig 7A are isocaloric, consuming an average of 2000 kcal/day. This is to some extent the case also in Fig 7B, comparing LCHF and HCLF (see Discussion: Limitations). The average behaviors of these diets are plotted in Fig 7C, for key variables. Finally, we also analyze the impact of ingestion speed and a change in weight. The model predicts that a high ingestion speed leads to a faster response, but there is no noticeable impact on amplitude of the key variables (Fig 7D). In the simulations, a lower body weight implies a higher insulin response, because the meal size is higher per body weight. This change in insulin dynamics implies higher glycogen levels, and lower EGP, which in turn inhibits gluconeogenesis. Note that our model does not have an insulin resistance term that depends on body weight, and that all other model parameters are kept the same in Fig 7E. In a real weight-loss study, it is likely that there will be an impact on insulin resistance. However, our simulations illustrate how one can isolate the function of the different components, in a way that is not possible *in vivo*.

## Discussion

We present a new digital twin that integrates many different studies - which all contain different, non-connected, and complementary information about human metabolism - into one useful, quantitative, big picture. The basis for this new twin technology is a multi-timescale, mechanistic, mathematical model (Fig 1A,D, S1), which describes the diet response of both the intracellular liver metabolism and the organ-organ crosstalk. The newly added and improved features in the model are centred on protein dynamics, intracellular liver metabolism, and the long-term regulations involving glycogen (Fig 1C, blue; Fig S1, blue). Two qualitative improvements (Fig 2A) illustrate the functionality of the new model mechanisms, and they are quantitatively tested in our new clinical study (Fig 6): i) glucose decreases over long-term fasting, since EGP comes from finite intracellular sources (Fig 2Aiii,2B,6A), ii) protein metabolism differs between a fed and a fasting state, since the energy-status in the body regulates whether protein should be used for gluconeogenesis or anabolic processes (Fig 2A,6B–C). The quantitative performance of the model is shown by *χ*^2^-test, which the model passes both for the training data (Fig 3), and the independent validation data (Fig 4–6). Finally, we demonstrate the potential usefulness of our digital twin tool, by simulating the impact that a variety of different diets is expected to have on key variables, such as fasting glucose, glucose AUC, fasting insulin, glycogen levels, which thereby provides a basis for understanding the consequences of these different diets (Fig 7). Many of these data and predictions (Fig 3D,4B,6,7) are subject-specific. We believe that this twin could become useful, not only for personalized diet-design, but also to improve patient understanding, motivation, compliance, and outcome (Shamanna et al., 2020).

### We can use the tool to estimate non-measured variables in existing and future studies, and thereby deepen our understanding of physiological processes and regulations

Our new digital twin tool can predict the response of both measured and non-measured variables. Measured variables correspond to known data and can be used to assess the model’s descriptive (Fig 3) or predictive (Fig 4–6) ability, depending on if the data is used for training (Fig 3) or testing (Fig 2B,E,4–6) the model, respectively. Non-measured variables in a specific study setting show the behavior of the model in variables for which we have no data. In other words, non-measured variables are predictions with the model. Examples of such non-measured variables are shown in Fig 2D, where we see gluconeogenesis, hepatic glycogen, and renal EGP dynamics; in Fig 2E, where we see the relative contributions of the different organs to glucose and insulin clearance and production; and in Fig 7, where we predict the response to different diets. Such predictions are possible and interesting to consider because our predictive tool already has demonstrated its ability to predict new variables (Fig 4–6), and because it utilizes information from other similar studies, where these variables have been measured. Finally, the model structure as such also provides valuable information, since it includes a sufficient set of mechanisms that can describe all these studied dynamics. For instance, in this model, the only long-term regulation variable is glycogen, which together with the short-term insulin regulation provide all the regulations included in the model. This means that, at least for these meal responses, the high complexity in metabolic regulation seen using gene-expression, bioinformatics, and other omics technologies, in the end, boils down to one major short-term and one major long-term regulation, which correlates with insulin and glycogen levels, respectively.

### The predictive ability of the model has been tested both quantitatively and qualitatively in a variety of ways

The quantitative assessments of the model predictions are shown in Figures 4–6, where the data (error bars) and simulations (lines and areas) are plotted together. Figure 4 shows such comparisons on an individual level, in a study where the subjects are fasting 60+ h. This is impressive because the model has only been trained on up to 23 h fasting, and only on average population data. Figure 5 shows validation tests using mixed-meal and OGTT studies, and examines both plasma glucose, plasma insulin, hepatic glycogen, and EGP dynamics (Firth et al., 1986, Taylor et al., 1996). Figure 6 shows our new clinical study, which validates the qualitatively new prediction (Fig 2A) that an OPTT gives a negligible response in plasma glucose levels before fasting, but a 1-2 mM response after 48 h fasting. This last prediction is important because this combination of OPTT and fasting is a fundamentally new type of observation, not present in any of the existing data used for either training or validation. The prediction is also reasonable, given basic biochemical knowledge: when you have fasted long enough, the liver metabolism switches to produce glucose from proteins, and when new proteins become available, hepatic glucose production goes up. However, while this strengthens the intuition behind why the prediction is reasonable, a quantitative model is needed to be able to predict how long one must wait, and how big the OPTT must be to produce a plasma glucose response of a certain magnitude. Note also that the organ proportions for glucose uptake in fasting conditions (Fig 2E,i), EGP production (Fig 2E, ii), and insulin degradation (Fig 2E, iii) all are in good agreement with data (Gerich, 2010, Mather and Pollock, 2011, Najjar and Perdomo, 2019). Finally, all the quantitative assessments above are clear both from a visual comparison between simulations and data, as well as from formal and statistical analysis using a *χ*^2^-test.

### Our new tool quantitatively predicts short- and intermediate-term responses to different diets

To illustrate an additional usage of our digital twin technology, and also to provide additional validations of model predictions, we simulated short-(hours) and intermediate-term (up to four weeks) responses to various dietary and fasting strategies (Fig 7). To isolate the impact of the dietary strategy as such, all compared options are isocaloric, seen over a week, to the extent that the model can obtain this. Using our analysis, we could for example compare the effect of diets with higher (SFM) and lower (IF and 5:2) meal frequencies (Fig 7A). We found that higher meal frequencies lead to clear reductions in mean plasma insulin levels, but to small changes in mean plasma glucose, which is in agreement with several previous studies (Holmstrup et al., 2010, Paoli et al., 2019, Taylor and Garrow, 2001). We can also predict additional impacts, not measured in those previous studies, for example that higher meal frequency diets give higher mean hepatic glycogen levels, and that less of the ingested protein is converted into glucose. This prediction is reasonable, since less frequent meals lead to more extended periods of fasting when glycogen is depleted, and gluconeogenesis is turned on. Similarly, diets containing high carbohydrates and lower protein (HCLF) resulted in higher hepatic glycogen, plasma insulin, and plasma glucose levels, which agrees with existing clinical studies (Ahmed et al., 2020, Shin et al., 2009). Given our results, one can also argue that diets such as LCHF, which lowers both average insulin and glucose levels, may be beneficial for treating diabetes and cardiovascular diseases, and short-term clinical studies indicate that LCHF does indeed lower fat-levels in both liver and blood (Mardinoglu et al., 2018). However, such clinical predictions should be done with great care, since several critical factors still are missing from the model. One such important missing factor is crosstalk between lipid and glucose/protein metabolism, which is currently not available in our, or in any other similar data-driven model. Furthermore, the model does not consider important long-term effects of actually prescribing the diets described here in clinical practice, since real clinical effects also involve impact on adherence and relapse risk, impact on appetite, etc, which are not included in the model. It may be because of such complicating factors, be that long-term studies of different diets show few clear results regarding which diet is best, and that effects of more well-controlled shorter studies show more clear effects. In any case, to be able to simulate the short- and intermediate term impact of following a diet may be useful to explain to a patient why we believe that the prescribed diet may be beneficial, and such simulations could increase patient understanding and motivation to adhere to the diet.

### Limitations, strengths, and key assumptions in our model compared to other available meal simulations models

Meal simulation models have been developed since the 1970s (Bergman et al., 1979, Bergman et al., 1981), but even though much developments has been done since then, there does not exist a previous model that can incorporate and predict/describe the data in the studies we have utilized herein (Fig 1E). An important step forward was the Dalla Man model (Dalla Man et al., 2007), which made use of triple-tracer data to construct a more physiological model, based on measurements estimating fluxes such as EGP, glucose uptake, and rate of appearance from the intestines. Our model can describe all of those data (Fig 2C, grey). Later models by us (Nyman et al., 2011, Herrgårdh et al., 2021) and others (Maas et al., 2015, Kurata, 2021) have produced more refined versions, which also describe glucose uptake subdivided into the different organs. Our model maintains this capacity and also describes organ-specific insulin clearance and EGP contributions (Fig 2E). Some other related previous models have instead focused on the detailed metabolism of the liver (König et al., 2012, Berndt et al., 2018, Ashworth et al., 2016), but such models have had no realistic meal response part. There are also various detailed models which have had little to no quantitative agreement with data, especially independent validation data (Kurata, 2021, Xu et al., 2011). Also, neither of these meal response models have described more long-term changes, such as build-up or breakdown of glycogen over days. There are more long-term models involving glycogen (Hall, 2009), but these models do not describe meal responses.

There are many shortcomings and assumptions in the model, many of which are due to a lack of the needed clinical data. For instance, the model does not include metabolism and crosstalk with lipids. This is a shortcoming in the context of simulation of LCHF, since the ingested fat is just assumed to be consumed, with no consequence for other dynamics. This shortcoming also means that an extension of these dynamics to more long-term dynamics, involving factors such as weight change, are not yet possible. There is currently no other model that describes such a crosstalk, and to develop such a glucose-protein-fat metabolism model is an important next step for the field, and it will bring many of the benefits hinted herein to an even more realized potential. Also, much of the intracellular metabolic reactions are highly simplified. To add such more detailed intracellular models, which also incorporate *in vitro* experimental data, is another important future research direction. We have previously done such multi-level expansions for insulin signalling in the adipose tissue (Brannmark et al., 2013, Herrgårdh et al., 2021), and those more detailed adipose tissue models can be included also in this model.

Despite these and other shortcomings with the presented model, it is showing the way towards a more all-encompassing meal simulation paradigm, that goes beyond glucose-insulin dynamics, towards more complex meal responses and long-term dynamics, which lays the basis for a new type of physiological digital twin technology. Such digital twin technologies have previously been developed for other processes, such as blood pressure and flow (Corral-Acero et al., 2020, Golse et al., 2021). For complex meal responses involving a variety of metabolites, only machine-learning models have previously been available (Shamanna et al., 2020). Using our new type of physiological models, we can go beyond such ‘black-box’ machine learning models to explainable AI. This opens the door to conveying medical information, which in turn could lead to increased patient motivation and compliance to dietary prescriptions, which in turn could help prevent metabolic and cardiovascular diseases.

## Material and method

### Mathematical modelling

The general mathematical modelling methodology used is similar to many of our previous papers (Nyman et al., 2014, Rajan et al., 2016), and below a short method summary is outlined. The mathematical analysis, model simulation, numerical optimization of model parameters, was carried out using MATLAB 2019b and the IQM toolbox (Schmidt and Jirstrand, 2006). All code is available at our GitHub repository, see: https://github.com/Orrestam/Digital-twin-predicting-diet-response-before-and-after-long-term-fasting.

The model is based on *ordinary differential equations* (ODEs) with the general form:

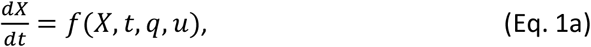

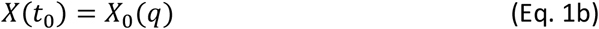

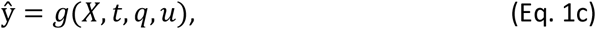

where *X* denotes a vector of state variables usually corresponding to concentrations of given system components; the functions *f* and *g* are non-linear smooth functions; *q* is a vector of model parameters (rate constants, scaling constants, etc.); *u* is the input signal corresponding to the meals ingested, as specified in the experimental data; *X*(*t*_0_) denotes the initial condition value *X*_0_(*q*), which are dependent on the model parameters *q*; and 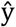 is the simulated model output. Parameter estimation was done by quantifying the model performance, using the model output 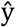 to calculate the traditional weighed least squares cost function written as

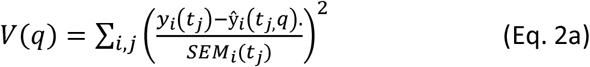

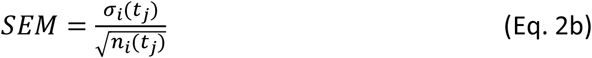

where *y*_*i*_(*t*_*j*_) denotes the measured datapoint, for a specific experimental setup and variable *i*, at time point *j*; where 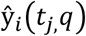 denotes the corresponding simulation value; and SEM is the standard error of the mean, which is the sample standard deviation, *σ*_*i*_(*t*_*j*_), divided with the square root of the number of repeats, *n*_*i*_(*t*_*j*_) at each time point. The value of the cost function, *V*(*q*), is then minimized by tuning the values of the parameters, typically referred to as parameter estimation. The parameter estimation was done using the particle swarm optimization algorithm from the MATLAB global optimization toolbox.

In order to evaluate the new model, we performed a *χ*^2^-test for the size of the residuals, with the null hypothesis that the experimental data have been generated by the model, and that the experimental noise is additive and normally distributed (Cedersund and Roll, 2009). In practice, the cost function value was compared to a *χ*^2^ test statistic, 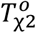. The test statistic value is given by the inverse *χ*^2^ cumulative density function,

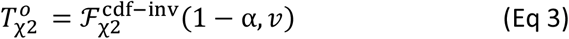

where 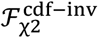 is the inverse density function; α is the significance level (α = 0.05, was used); and *v* is the degrees of freedom, which was equal to the number of data points in the training dataset (273 in total, all timepoints over all experiments). In practice, the model is rejected if the model cost is larger than the *χ*^2^-threshold 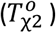.

### Qualitative constraints to the optimization

To avoid unphysiological behaviours, we also added several *ad hoc* requirements to the optimization. These requirements, if not fully filled, would increase the cost function (Eq. 2a) with a constant term, which is present during the optimization, but not during the *χ*^2^-test. These *ad hoc* requirements were based on e.g. the data in Fig 2E:

- No negative reaction flows, state values, or variables.
- No plasma insulin above 3000 pmol/L.
- Renal glucose production is not permitted to exceed 40% of the total glucose production (Mather and Pollock, 2011).
- Insulin degradation in liver is not permitted to decrease below 40% (Najjar and Perdomo, 2019).
- Glucose uptake ratios to the different organs, both during postprandial and overnight fasting, should be reasonably proportionate (Fig 2E,i) (Gerich, 2010).
- Gluconeogenesis contribution to the endogenous glucose production (EGP) is not allowed to exceed 100%.

### Seven existing clinical studies used for estimation and validation

A total of seven existing clinical studies (Krssak et al., 2004, Rothman Douglas et al., 1991, Magnusson et al., 1992, Firth et al., 1986, Lerche et al., 2009, Taylor et al., 1996, Dalla Man et al., 2007) were collected to train (Fig 3) and validate (Fig 4–5) the model. The studies used are presented in Fig 1C. Basic features of these studies are described in the legends to Figures 3–5, and more details are summarized in Fig S4. For a full description of the methodologies used, we refer to the individual papers. Most of the data used was available in the Supplementary material (König et al., 2012), and the remaining data were extracted from the figures in the papers. In data of Lerche *et al*. (2009) (Lerche et al., 2009) the SEM of insulin in two datapoints were particularly small and to avoid a too high weight on those datapoints, their SEM was thus replaced with the mean SEM during the fasting intervention.

### New clinical study: OPTT before and after 2 days of fasting

While there are many highly informative studies available in the literature, we found a critical lack of data for certain combinations of diets, meal sizes, and fasting schedules. Therefore, a new study was conducted (Fig 6A), to provide some critical information needed to train the model, and to allow for a completely independent validation of critical predictions with the model (Fig 2A,iii, Fig 6). Since our model is a digital twin technology, with individual models for each subject, it is more important that N is high for the number of data points, and an N=1 study is in principle sufficient regarding the number of subjects. For this reason, we performed a study with only 3 non-diabetic subjects (N=3, ages= [24,29,42] years, weights = [80,87,85] kg, heights = [1.78,1.79, 1.85] m). To remedy the need for a high time-resolution, we combined a wearable continuous glucose monitor (CGM) (Abbott, *FreeStyle Libre 2*), with a regular sampling of blood glucose values using the associated Abbott kit, to allow for calibration of the CGM values. The CGM sampled every 15 minutes throughout the entire 14-days study, and during the critical OPTT response, we collected blood samples every 2-4 minutes. During the first 3 days of the study, normal non-controlled diet was followed, but some meals were carefully measured, to allow for calibrations of the digital twins. Importantly, at least one of these meals was large, ~1000 kcal, since such large meal responses are lacking from all other clinical studies included in the analyzed data. For this reason, one such large meal response was included in the estimation data (Fig 3D). At the end of day 3, a regular dinner was ingested at 19.00, and the first OPTT was taken at 23.00 (132.5 kcal, 25.55 g proteins, 2.6 g carbohydrates). After this, a 48 h fast followed (only water and black tea allowed), and at 23.00 on day 5, a new OPTT was taken, using the same protocol. After this second OPTT, CGM values were collected for another 9 days, until the sensor expired. The study was conducted with the approval of the local ethics board, DNR-2021-02668.

## Supporting information

Complete model

## Author contributions

OS has done the model development and analysis. Primary supervision and original conceptions were done by GC. Additional supervision and inputs on parts of the modelling come from CS. PG contributed to model formulations and descriptions, and PL and ME contributed with clinical perspectives. The manuscript was primarily written by GC, with primary input from OS and CS. All authors contributed to the text and discussions, and approved the final manuscript.

## References

Ahmed, S. R., Bellamkonda, S., Zilbermint, M., Wang, J. & Kalyani, R. R. 2020. Effects of the low carbohydrate, high fat diet on glycemic control and body weight in patients with type 2 diabetes: experience from a community-based cohort. BMJ open diabetes research & care, 8, e000980.

Ajala, O., English, P. & Pinkney, J. 2013. Systematic review and meta-analysis of different dietary approaches to the management of type 2 diabetes. The American Journal of Clinical Nutrition, 97, 505–516.

Ashworth, W. B., Davies, N. A. & Bogle, I. D. L. 2016. A Computational Model of Hepatic Energy Metabolism: Understanding Zonated Damage and Steatosis in NAFLD. PLOS Computational Biology, 12, e1005105.

Bergman, R. N., Ider, Y. Z., Bowden, C. R. & Cobelli, C. 1979. Quantitative estimation of insulin sensitivity. American Journal of Physiology-Endocrinology and Metabolism, 236, E667.

Bergman, R. N., Phillips, L. S. & Cobelli, C. 1981. Physiologic evaluation of factors controlling glucose tolerance in man: measurement of insulin sensitivity and beta-cell glucose sensitivity from the response to intravenous glucose. The Journal of clinical investigation, 68, 1456–1467.

Berndt, N., Bulik, S., Wallach, I., Wunsch, T., Konig, M., Stockmann, M., Meierhofer, D. & Holzhutter, H. G. 2018. Hepatokin1 is a biochemistry-based model of liver metabolism for applications in medicine and pharmacology. Nat Commun, 9, 2386.

Brannmark, C., Nyman, E., Fagerholm, S., Bergenholm, L., Ekstrand, E. M., Cedersund, G. & Stralfors, P. 2013. Insulin signaling in type 2 diabetes: experimental and modeling analyses reveal mechanisms of insulin resistance in human adipocytes. J Biol Chem, 288, 9867–80.

Cedersund, G. & Roll, J. 2009. Systems biology: model based evaluation and comparison of potential explanations for given biological data. FEBS journal, 276, 903–922.

Chawla, S., Tessarolo Silva, F., Amaral Medeiros, S., Mekary, R. A. & Radenkovic, D. 2020. The Effect of Low-Fat and Low-Carbohydrate Diets on Weight Loss and Lipid Levels: A Systematic Review and Meta-Analysis. Nutrients, 12, 3774.

Cobelli, C., Pacini, G., Toffolo, G. & Sacca, L. 1986. Estimation of insulin sensitivity and glucose clearance from minimal model: new insights from labeled IVGTT. American Journal of Physiology-Endocrinology and Metabolism, 250, E591–E598.

Cobelli, C. & Thomaseth, K. 1987. The minimal model of glucose disappearance: optimal input studies. Mathematical Biosciences, 83, 127–155.

Corral-Acero, J., Margara, F., Marciniak, M., Rodero, C., Loncaric, F., Feng, Y., Gilbert, A., Fernandes, J. F., Bukhari, H. A., Wajdan, A., Martinez, M. V., Santos, M. S., Shamohammdi, M., Luo, H., Westphal, P., Leeson, P., Diachille, P., Gurev, V., Mayr, M., Geris, L., Pathmanathan, P., Morrison, T., Cornelussen, R., Prinzen, F., Delhaas, T., Doltra, A., Sitges, M., Vigmond, E. J., Zacur, E., Grau, V., Rodriguez, B., Remme, E. W., Niederer, S., Mortier, P., Mcleod, K., Potse, M., Pueyo, E., Bueno-Orovio, A. & Lamata, P. 2020. The ‘Digital Twin’ to enable the vision of precision cardiology. European heart journal, 41, 4556–4564.

Dalla Man, C., Rizza, R. A. & Cobelli, C. 2007. Meal simulation model of the glucose-insulin system. IEEE Trans Biomed Eng, 54, 1740–9.

Dashti, H. S. & Mogensen, K. M. 2017. Recommending Small, Frequent Meals in the Clinical Care of Adults: A Review of the Evidence and Important Considerations. Nutrition in Clinical Practice, 32, 365–377.

Duckworth, W. C., Bennett, R. G. & Hamel, F. G. 1998. Insulin Degradation: Progress and Potential*. Endocrine Reviews, 19, 608–624.

Firth, R. G., Bell, P. M., Marsh, H. M., Hansen, I. & Rizza, R. A. 1986. Postprandial hyperglycemia in patients with noninsulin-dependent diabetes mellitus. Role of hepatic and extrahepatic tissues. The Journal of clinical investigation, 77, 1525–1532.

Ganesan, K., Habboush, Y. & Sultan, S. 2018. Intermittent Fasting: The Choice for a Healthier Lifestyle. Cureus, 10, e2947–e2947.

García-Pérez, L.-E., Alvarez, M., Dilla, T., Gil-GuillÉn, V. & Orozco-BeltrÁn, D. 2013. Adherence to therapies in patients with type 2 diabetes. Diabetes therapy: research, treatment and education of diabetes and related disorders, 4, 175–194.

Ge, L., Sadeghirad, B., Ball, G. D. C., Da Costa, B. R., Hitchcock, C. L., Svendrovski, A., Kiflen, R., Quadri, K., Kwon, H. Y., Karamouzian, M., Adams-Webber, T., Ahmed, W., Damanhoury, S., Zeraatkar, D., Nikolakopoulou, A., Tsuyuki, R. T., Tian, J., Yang, K., Guyatt, G. H. & Johnston, B. C. 2020. Comparison of dietary macronutrient patterns of 14 popular named dietary programmes for weight and cardiovascular risk factor reduction in adults: systematic review and network meta-analysis of randomised trials. BMJ, 369, m696.

Gerich, J. E. 2010. Role of the kidney in normal glucose homeostasis and in the hyperglycaemia of diabetes mellitus: therapeutic implications. Diabetic medicine: a journal of the British Diabetic Association, 27, 136–142.

Gibson, A. A. & Sainsbury, A. 2017. Strategies to Improve Adherence to Dietary Weight Loss Interventions in Research and Real-World Settings. Behavioral sciences (Basel, Switzerland), 7, 44.

Golse, N., Joly, F., Combari, P., Lewin, M., Nicolas, Q., Audebert, C., Samuel, D., Allard, M.-A., Sa Cunha, A., Castaing, D., Cherqui, D., Adam, R., Vibert, E. & Vignon-Clementel, E. 2021. Predicting the risk of post-hepatectomy portal hypertension using a digital twin: A clinical proof of concept. Journal of Hepatology, 74, 661–669.

Hall, K. D. 2009. Predicting metabolic adaptation, body weight change, and energy intake in humans. American Journal of Physiology-Endocrinology and Metabolism, 298, E449–E466.

Herrgårdh, T., Li, H., Nyman, E. & Cedersund, G. 2021. An Updated Organ-Based Multi-Level Model for Glucose Homeostasis: Organ Distributions, Timing, and Impact of Blood Flow. Frontiers in physiology, 12, 619254–619254.

Holmstrup, M. E., Owens, C. M., Fairchild, T. J. & Kanaley, J. A. 2010. Effect of meal frequency on glucose and insulin excursions over the course of a day. e-SPEN, the European e-Journal of Clinical Nutrition and Metabolism, 5, e277–e280.

Jaworski, M., Panczyk, M., Cedro, M. & Kucharska, A. 2018. Adherence to dietary recommendations in diabetes mellitus: disease acceptance as a potential mediator. Patient preference and adherence, 12, 163–174.

Kovatchev, B. P., Breton, M., Man, C. D. & Cobelli, C. 2009. In silico preclinical trials: a proof of concept in closed-loop control of type 1 diabetes. Journal of diabetes science and technology, 3, 44–55.

Krssak, M., Brehm, A., Bernroider, E., Anderwald, C., Nowotny, P., Man, C. D., Cobelli, C., Cline, G. W., Shulman, G. I., WaldhÄusl, W. & Roden, M. 2004. Alterations in Postprandial Hepatic Glycogen Metabolism in Type 2 Diabetes. Diabetes, 53, 3048.

Kurata, H. 2021. Virtual metabolic human dynamic model for pathological analysis and therapy design for diabetes. iScience, 24, 102101–102101.

KÖnig, M., Bulik, S. & HolzhÜtter, H.-G. 2012. Quantifying the Contribution of the Liver to Glucose Homeostasis: A Detailed Kinetic Model of Human Hepatic Glucose Metabolism. PLOS Computational Biology, 8, e1002577.

Lerche, S., Soendergaard, L., Rungby, J., Moeller, N., Holst, J. J., Schmitz, O. E. & Brock, B. 2009. No increased risk of hypoglycaemic episodes during 48 h of subcutaneous glucagon-like-peptide-1 administration in fasting healthy subjects. Clinical Endocrinology, 71, 500–506.

Maas, A. H., Rozendaal, Y. J. W., Van Pul, C., Hilbers, P. A. J., Cottaar, W. J., Haak, H. R. & Van Riel, N. A. W. 2015. A physiology-based model describing heterogeneity in glucose metabolism: the core of the Eindhoven Diabetes Education Simulator (E-DES). Journal of diabetes science and technology, 9, 282–292.

Magnusson, I., Rothman, D. L., Katz, L. D., Shulman, R. G. & Shulman, G. I. 1992. Increased rate of gluconeogenesis in type II diabetes mellitus. A 13C nuclear magnetic resonance study. The Journal of clinical investigation, 90, 1323–1327.

Mardinoglu, A., Wu, H., Bjornson, E., Zhang, C., Hakkarainen, A., Rasanen, S. M., Lee, S., Mancina, R. M., Bergentall, M., Pietilainen, K. H., Soderlund, S., Matikainen, N., Stahlman, M., Bergh, P. O., Adiels, M., Piening, B. D., Graner, M., Lundbom, N., Williams, K. J., Romeo, S., Nielsen, J., Snyder, M., Uhlen, M., Bergstrom, G., Perkins, R., Marschall, H. U., Backhed, F., Taskinen, M. R. & Boren, J. 2018. An Integrated Understanding of the Rapid Metabolic Benefits of a Carbohydrate-Restricted Diet on Hepatic Steatosis in Humans. Cell Metab, 27, 559–571 e5.

Mather, A. & Pollock, C. 2011. Glucose handling by the kidney. Kidney International, 79, S1–S6.

Najjar, S. M. & Perdomo, G. 2019. Hepatic Insulin Clearance: Mechanism and Physiology. Physiology (Bethesda, Md.), 34, 198–215.

Noakes, T. D. & Windt, J. 2017. Evidence that supports the prescription of low-carbohydrate high-fat diets: a narrative review. British Journal of Sports Medicine, 51, 133.

Nyman, E., Brannmark, C., Palmer, R., Brugard, J., Nystrom, F. H., Stralfors, P. & Cedersund, G. 2011. A hierarchical whole-body modeling approach elucidates the link between in Vitro insulin signaling and in Vivo glucose homeostasis. J Biol Chem, 286, 26028–41.

Nyman, E., Rajan, M. R., Fagerholm, S., Brännmark, C., Cedersund, G. & Strålfors, P. 2014. A single mechanism can explain network-wide insulin resistance in adipocytes from obese patients with type 2 diabetes. The Journal of biological chemistry, 289, 33215–33230.

Nyman, E., Rozendaal, Y. J., Helmlinger, G., Hamren, B., Kjellsson, M. C., Stralfors, P., Van Riel, N. A., Gennemark, P. & Cedersund, G. 2016. Requirements for multi-level systems pharmacology models to reach end-usage: the case of type 2 diabetes. Interface Focus, 6, 20150075.

Paoli, A., Tinsley, G., Bianco, A. & Moro, T. 2019. The Influence of Meal Frequency and Timing on Health in Humans: The Role of Fasting. Nutrients, 11, 719.

Rajan, M. R., Nyman, E., KjøLhede, P., Cedersund, G. & Strålfors, P. 2016. Systems-wide Experimental and Modeling Analysis of Insulin Signaling through Forkhead Box Protein O1 (FOXO1) in Human Adipocytes, Normally and in Type 2 Diabetes. Journal of Biological Chemistry, 291, 15806–15819.

Rothman Douglas, L., Magnusson, I., Katz Lee, D., Shulman Robert, G. & Shulman Gerald, I. 1991. Quantitation of Hepatic Glycogenolysis And Gluconeogenesis in Fasting Humans With 13C NMR. Science, 254, 573–576.

Schmidt, H. & Jirstrand, M. 2006. Systems Biology Toolbox for MATLAB: a computational platform for research in systems biology. Bioinformatics, 22, 514–5.

Scholtens, E. L., Krebs, J. D., Corley, B. T. & Hall, R. M. 2020. Intermittent fasting 5:2 diet: What is the macronutrient and micronutrient intake and composition? Clinical Nutrition, 39, 3354–3360.

Schwartz, S. M., Wildenhaus, K., Bucher, A. & Byrd, B. 2020. Digital Twins and the Emerging Science of Self: Implications for Digital Health Experience Design and “Small” Data. Frontiers in Computer Science, 2.

Shamanna, P., Saboo, B., Damodharan, S., Mohammed, J., Mohamed, M., Poon, T., Kleinman, N. & Thajudeen, M. 2020. Reducing HbA1c in Type 2 Diabetes Using Digital Twin Technology-Enabled Precision Nutrition: A Retrospective Analysis. Diabetes Therapy, 11, 2703–2714.

Shin, Y., Park, S. & Choue, R. 2009. Comparison of time course changes in blood glucose, insulin and lipids between high carbohydrate and high fat meals in healthy young women. Nutrition research and practice, 3, 128–133.

Sips, F. L. P., Nyman, E., Adiels, M., Hilbers, P. A. J., Strålfors, P., Van Riel, N. A. W. & Cedersund, G. 2015. Model-Based Quantification of the Systemic Interplay between Glucose and Fatty Acids in the Postprandial State. PLOS ONE, 10, e0135665.

Sylvetsky, A. C., Edelstein, S. L., Walford, G., Boyko, E. J., Horton, E. S., Ibebuogu, U. N., Knowler, W. C., Montez, M. G., Temprosa, M., Hoskin, M., Rother, K. I., Delahanty, L. M. & Diabetes Prevention Program Research, G. 2017. A High-Carbohydrate, High-Fiber, Low-Fat Diet Results in Weight Loss among Adults at High Risk of Type 2 Diabetes. The Journal of nutrition, 147, 2060–2066.

Taylor, M. A. & Garrow, J. S. 2001. Compared with nibbling, neither gorging nor a morning fast affect short-term energy balance in obese patients in a chamber calorimeter. International Journal of Obesity, 25, 519–528.

Taylor, R., Magnusson, I., Rothman, D. L., Cline, G. W., Caumo, A., Cobelli, C. & Shulman, G. I. 1996. Direct assessment of liver glycogen storage by 13C nuclear magnetic resonance spectroscopy and regulation of glucose homeostasis after a mixed meal in normal subjects. The Journal of clinical investigation, 97, 126–132.

Xu, K., Morgan, K. T., Todd Gehris, A., Elston, T. C. & Gomez, S. M. 2011. A Whole-Body Model for Glycogen Regulation Reveals a Critical Role for Substrate Cycling in Maintaining Blood Glucose Homeostasis. PLOS Computational Biology, 7, e1002272.

