## Supplementary material for "Digital twin predicting diet response before and after long-term fasting": Complete model

a) Department of Biomedical Engineering, IMT, Linköping University, Linköping, Sweden
b) Center for Medical Image Science and Visualization, Linköping University, Linköping, Sweden
c) Department of Health, Medicine, and Caring Sciences, Linköping University, Linköping, Sweden
d) Division of Medical Radiation Physics, at the Department of Health, Medicine and Caring Sciences, Linköping University, Linköping, Sweden
e) Drug Metabolism and Pharma
d) Drug Metabolism and Pharmacokinetics, Research and Early Development, Cardiovascular, Renal and Metabolism (CVRM), BioPharmaceuticals R&D, AstraZeneca, Gothenburg, Sweden

### Introduction

The proposed model extends a previously published short timescale model [1] and now serves on a multi-timescale with the ability to simulate metabolic flexibility during both short time periods, such as a meal, and slightly longer timer periods, up to a few days. This extension to longer timescales has required us to add a new organ to the model, the liver, and to describe some of its metabolic processes, such as glycogen synthesis and breakdown, protein synthesis and breakdown, digestion of meals containing protein, and long-term regulation of the corresponding fluxes. An overview of the old and the new model is given in Figure 1, where the old model is color-coded red and new parts are color-coded blue.

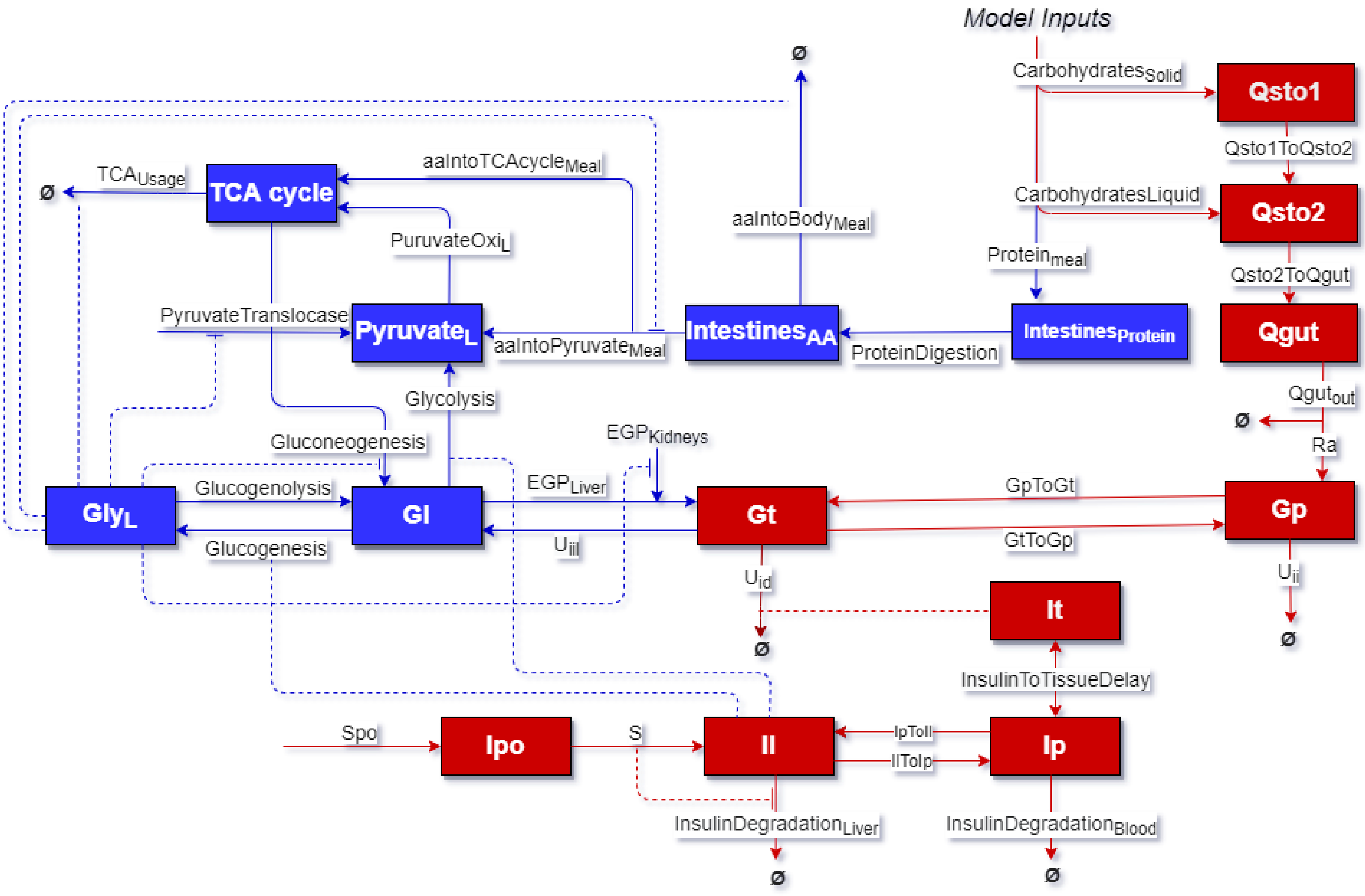

*Figure S1. Illustration of the complete model. Rectangles represent states, text with no squares are model reactions, and flows into Ø are flows leading out of model. Red colour represents the old model and blue the new. Dotted lines represent positive dependency regulations and dotted lines ending with a perpendicular head represent inhibitions.*

### Model structure

The model consists of i) states, ii) parameters, iii) events, and iv) the ordinary differential equations, and their constituent reactions. We will go through these one by one.

### Model states

The model states are dynamic quantities which were illustrated in Figure 1 as rectangles. The states are listed in Table S1.

*Table S1, List of model states, its corresponding units, and a short description of what it represents. Initial value of simulations is represented as* $\theta_0$.

| Category | Name | Unit | Description | $\theta_0$ |
|---|---|---|---|---|
| Stomach | **Qsto1** | mg | Polysaccharides in stomach | 1e-50 |
| | **Qsto2** | mg | Maltose/sugar/lactose in stomach | 0 |
| | **Qgut** | mg | Carbohydrates in gut | 0 |
| | **$Intestine_{Protein}$** | mg/kg | Protein in intestines | 1e-50 |
| | **$Intestine_{aa}$** | mg/kg | Amino acids in intestines | 0 |
| Blood | **Gp** | mg/kg | Glucose in plasma | 80 |
| | **Ip** | pmol/kg | Insulin in plasma | 1 |
| | **Ipo** | pmol/kg | Insulin in portal vein | 0 |
| Tissue | **Gt** | mg/kg | Glucose in tissue | 50 |
| Liver | **Gl** | mg/kg | Glucose in liver | 20 |
| | **$Gly_L$** | mg/kg | Hepatic glycogen | 1500 |
| | **$Pyruvate_L$** | mg/kg | Pyruvate in liver | 50 |
| | **$TCAcycle_L$** | mg/kg | Sum of TCA cycle components in liver | 50 |
| | **Il** | pmol/kg | Insulin in liver | 1 |
| Global | **InsulinStabilization** | pmol/kg | Part of insulin production calculations | 0 |
| | **InsulinToTissueDelay** | pmol/L | Insulin response in tissue | 0 |

To ensure that the model is in a realistic steady state and in agreement with data at time point 0, the model simulation starts 5 days prior to day 0. Because all data has different measured initial variables the steady state simulations of 5 days contain a standardized diet that contains different amounts of calories, optimized to each dataset. For this reason, the initial values of all states in a particular simulation at day 0 will generally deviate from the initial parameters of Table S1.

### Model parameters

Model parameters are usually held constant throughout the simulation. An exception to this is when a parameter is changed by a model event. The values of the parameters may be specific to an individual, e.g. disease state, weight, and age. Because of this, the model can simulate individuals of different genders, weight and metabolic conditions such as insulin resistance and faster metabolism. The parameters of the model are given in Table S2.

*Table S2, List of parameters and the starting guess, **θ₀**, of each parameter in the parameter estimation. $\theta_{0\_Healthy}$ is the parameter starting guess for healthy populations and $\theta_{0\_T2DM}$ is for diabatic populations. The categorization observed in Table S2 is based on the location of corresponding physiological mechanism, and parameters of flows between categories are specified by the category of the start of the flow, e.g. the parameter representing glucose diffusion between blood and tissue, $k_1$, is specified in the blood category.*

| Category | Name | Unit | Description | $\theta_{0\_Healthy}$ | $\theta_{0\_T2DM}$ |
|---|---|---|---|---|---|
| Stomach | $k_{gri}$ | 1/min | Breakdown of polysaccharides to monosaccharides | 0,064 | 0,045 |
| | $k_{min}$ | 1/min | Minimum rate of gastric emptying | 0,009 | 0,007 |
| | $k_{max}$ | 1/min | Maximum rate of gastric emptying | 0,122 | 0,100 |
| | **b** | min | Time period of minimum rate of gastric emptying | 0,933 | 0,930 |
| | **d** | min | Time period of maximum rate of gastric emptying | 0,010 | 0,030 |
| | $k_{abs}$ | 1/min | Rate constant of flux from glucose in gut to glucose in blood | 0,028 | 0,020 |
| | $aaTransportion_K$ | 1/min | Transportation rate of amino acids from intestines | 0,001 | 0,001 |
| | **ProteinBreakdown** | 1/min | Breakdown rate of protein to amino acids in intestines | 0,013 | 0,020 |
| Blood | **f** | % | Percentage conversion of digested carbohydrate to glucose in blood | 0.900 | 0,957 |
| | $k_1$ | 1/min | Glucose diffusion constant between blood and tissue | 0,058 | 0,040 |
| | $U_{ii}$ | mg/kg/min | Insulin-independent glucose utilization from plasma | 0,544 | 0,724 |
| | $m_2$ | 1/min | Insulin diffusion constant between insulin in blood and liver | 1,038 | 0,455 |
| | $m_4$ | 1/min | Insulin degradation rate from blood | 0,150 | 0,040 |
| | **K** | % | Pancreatic responsivity to the glucose rate of change and unit conversion | 412,800 | 407,462 |
| | $S_b$ | pmol/kg/min | Basal insulin secretion | 0,0130 | 0,010 |
| | **gamma** | % | Pancreatic responsivity to glucose concentration | 447,915 | 298,334 |
| Tissue | $K_{m0}$ | 1/min | Rate of change of glucose utilization of muscle | 207,308 | 599,99 |
| | $V_{m0}$ | 1/min | Basal rate of glucose utilization of muscle | 3,511 | 1,100 |
| | $V_{mX}$ | Dimensionless | Insulin dependent rate of glucose utilization of muscle | 0,140 | 0,400 |
| | $K_{f0}$ | 1/min | Rate of change of glucose utilization of adipocytes | 290,242 | 469,308 |

| | | | | | |
|---|---|---|---|---|---|
| | **$V_{f0}$** | 1/min | Basal rate of glucose utilization of adipocytes | 0,048 | 2,999 |
| | **$V_{fX}$** | Dimensionless | Insulin dependent rate of glucose utilization of adipocytes | 0,145 | 0,190 |
| | **$EGPLiver_{diffusionMax}$** | 1/min | Maximum transportation rate of glucose out of liver | 3,962 | 2,590 |
| | **$EGPLiver_{diffusion0}$** | 1/min | Rate of change of glucose diffusion out of liver | 7,959 | 1,763 |
| | **$Uidl_{diffusionMax}$** | 1/min | Maximum transportation rate of glucose into the liver | 0,396 | 0,593 |
| | **$Uidl_{diffusion0}$** | 1/min | Transportation rate of glucose into liver up to maximum rate | 0,910 | 1,000 |
| | **$EGP_{KidneysK}$** | 1/min | Non dynamic kidney glucose production | 0,625 | 0,939 |
| | **$k_2$** | 1/min | Diffusion constant of glucose between tissue and blood | 0,238 | 0,547 |
| | **$It_{delayK}$** | Dimensionless | Delay between rate of appearance of insulin in plasma and insulin response in tissue. | 0,345 | 1,500 |
| | **$G_b$** | mg/kg | Basal glucose concentration | 27,703 | 34,638 |
| Liver | **$V_{glyBmax}$** | 1/min | Maximum rate of glycogenolysis | 4,732 | 6,000 |
| | **GlyB** | 1/min | Rate of glycogenolysis | 81,548 | 120,00 0 |
| | **$V_{glySmax}$** | 1/min | Maximum rate of glycogenesis | 4,736 | 1,721 |
| | **GlyS** | 1/min | Rate of glycogenesis | 0,801 | 0,400 |
| | **$TCAusage_K$** | 1/min | Rate of disappearance of amino acids in liver from the TCA cycle | 0,989 | 0,300 |
| | **$PyruvateTranslocase_K$** | 1/min | Rate of appearance of pyruvate from the body into the liver | 1,822 | 2,500 |
| | **$Gluconeogenesis_K$** | 1/min | Gluconeogenesis from the TCA cycle in the liver | 0,011 | 0,005 |
| | **$PyruvateOxi_K$** | 1/min | Oxidation of pyruvate to AcetylCoA in the liver | 0,028 | 0,050 |
| | **$Aminoprofile_K$** | % | Amino acid profile estimation from a meal | 0,762 | 0,358 |
| | **$Glycolysis_k$** | 1/min | Glycogen-independent rate constant of glycolysis | 0,101 | 0,456 |
| | **$Glycolysis_{EXP}$** | Dimensionless | Glycolysis dependency in relation to hepatic insulin response | 0,042 | 0,004 |
| | **$InsulinDep_{EXP}$** | Dimensionless | Exponential relation between usage of glucose and insulin | 1,404 | 1,001 |
| | **$m_1$** | 1/min | Insulin diffusion constant between insulin in the liver and glucose in plasma | 0,400 | 0,310 |

| | | | | | |
|---|---|---|---|---|---|
| | **$m_5$** | 1/min | Hepatic extraction of insulin dependent on the secretion of insulin between the portal vein and the liver | 0,128 | 0,082 |
| | **$m_6$** | 1/min | Basal hepatic extraction of insulin | 0,225 | 0,603 |
| Global | **alpha** | Dimensionless | Delay between glucose signal and insulin secretion | 4,430 | 3,127 |
| | **beta** | Dimensionless | Pancreatic response to hypoglycemia and hyperglycemia | 0,055 | 0,045 |
| | **InsulinLiverResponseK** | Dimensionless | Hepatic insulin response | 0,105 | 0,872 |
| | **$GlyDep_{Meal}$** | mg/kg | Amino acid transportation dependency on glycogen as an energy regulator | 464,126 | 413,868 |
| | **$GlyDep_{TCA}$** | mg/kg | Glucose utilization dependent on glycogen as an energy regulator | 254,494 | 347,22 |
| | **$GlyDep_{Gluconeogenesis}$** | mg/kg | Rate of gluconeogenesis dependent on glycogen as an energy regulator | 943,610 | 750,000 |
| | **$GlyDepInFlow_K$** | mg/kg | Flow into the model dependent on glycogen as an energy regulator | 44,171 | 50,017 |
| | **$GlyDepEXP_{Meal}$** | Dimensionless | Amount of protein used for gluconeogenesis dependent on glycogen as an energy regulator | 1,020 | 0,648 |
| | **$GlyDepEXP_{TCA}$** | Dimensionless | Glucose utilization dependent on glycogen as an energy regulator | 0,874 | 0,219 |
| | **$GlyDepEXP_{Gluconeogenesis}$** | Dimensionless | Rate of gluconeogenesis dependent on glycogen as an energy regulator | 1,237 | 0,900 |
| | **$GlyDepIn_{kEXP}$** | Dimensionless | Flow into the model dependent on glycogen as an energy regulator | 0,047 | 0,150 |
| | **BloodVolumeUncertainty** | % | Uncertainty in estimating total blood volume | 0.920 | 0.970 |
| | **BloodLiverUncerteinty** | % | Uncertainty in estimating blood volume in liver | 0.950 | 1.040 |

Because there are fundamental differences between populations and individuals, three pairs of parameters are allowed to vary between studies. Firstly, parameters **K** and **beta** allowed to vary with the same magnitude between populations to account for differences in production of insulin. Secondly, parameters **$m_5$**, **$m_4$** are allowed to vary with the same magnitude to account for differences in insulin degradiation. Lastly, parameters **$V_{mx}$**, **$V_{fx}$** are allowed to vary with the same magnitude to account for differences in insulin response in glucose utilization. Without these population specific parameter calibrations all populations metabolisms would behave exactly as a mean metabolism of all populations in estimation data and would thus highly limit the amount of usable validation and estimation data. All the datapoints used in testing the model prediction capabilities (Figures 4-6) are marked with a cross (X). This underlines that there is a need for our Digital twin tool to be calibrated to an individual or population before making specific predictions on for example a diet or meal.

Parameters can also be used to send information into the model, as a model input. By using such model inputs, the model can for example predict metabolic fluxes for a specific combination of meals eaten by an individual with a specific body composition. The model inputs are listed in Table S3.

*Table S3, List of model inputs.*

| Categorization | Name | Unit | Description |
|---|---|---|---|
| **Meal Information** | **$Meal_{Length}$** | min | Length of meal |
| | **$Meal_{Start}$** | min | Start time of meal |
| | **$Carbohydrate_{Amount}$** | g | Total amount of carbohydrates in meal |
| | **$Protein_{Amount}$** | g | Total amount of protein in meal |
| | **$MealSolid_{Boolean}$** | True or False | Boolean to determine if meal is in solid phase |
| | **$MealLiquid_{Boolean}$** | True or False | Boolean to determine if meal is in liquid phase |
| **Body Information** | **$Female_{Boolean}$** | True or False | Boolean to determine gender in estimation of blood volume |
| | **$Male_{Boolean}$** | True or False | Boolean to determine gender in estimation of blood volume |
| | **Height** | cm | Height of the person, used to estimate blood volume |
| | **$BW_{start}$** | kg | Body weight of the person |

### Model events

A specific stimulation is defined by events sent into the model. An event is a Boolean function that specifies a change when the variable **Time** becomes a pre-specified value. For example, a meal event starts when the **$Meal_{Boolean}$** is changed from a 0 to a 1. This happens when the vector **Time** is equal to the input **$Meal_{Start}$** and lasts until the vector **Time** is equal to **$Meal_{End}$**. **$Meal_{End}$** is a sum of model input **$Meal_{Length}$** and **$Meal_{Start}$**, see Equation 1.1.

$$\mathbf{Meal_{End}[min] = Meal_{Start}[min] + Meal_{Length}[min]} \qquad \text{Eq 1.1}$$

The phase of the meal is defined by the booleans **$MealSolid_{Boolean}$** and **$MealSolid_{Liquid}$**. During a meal event, the parameter **D** is set to $\mathbf{Meal_{Amount}}$, and is ultimately used to calculate the rate of the digestion of the meal in relation to its size. This meal digestion rate is further explained in Equation 5.1.1.

See Equation 1.1.1 – Equation 1.1.3 for meal events.

$$\mathbf{MealTurnOn = gt(Time, Meal_{Start}), Meal_{Boolean}, 1} \qquad \text{Eq 1.1.1}$$

$$\mathbf{StomachActivation = gt(time, Meal_{Start}), D, Meal_{Amount}} \qquad \text{Eq 1.1.2}$$

$$\mathbf{MealTurnOff = gt(Time, Meal_{End}), Meal_{Boolean}, 0} \qquad \text{Eq 1.1.3}$$

**$Meal_{Amount}$** in Equation 1.2. is the amount of carbohydrates in the meal. This is calculated through converting the model input $\mathbf{Carbohydrate_{Amount}}$ from [g] to [mg], see Equation 1.2.

$$\mathbf{Meal_{Amount}\ [mg] = 10^3 * Carbohydrate_{Amount}[g]} \qquad \text{Eq 1.2}$$

During a meal event a flow of macronutrients is sent into the model, see Equations 1.4.1 – Equation 1.4.3.

$$\mathbf{Carbohydrate_{Solid}[\frac{mg}{min}] = \frac{10^3 * CarbohydrateAmount[g]}{Meal_{Length}[min]} * MealSolid_{Boolean} * Meal_{Boolean}} \qquad \text{Eq 1.4.1}$$

$$\mathbf{Carbohydrate_{Liquid}}[\frac{\mathbf{mg}}{\mathbf{min}}] = \frac{\mathbf{10^3 * CarbohydrateAmount[g]}}{\mathbf{Meal_{Length}[min]}} * \mathbf{MealLiquid_{Boolean}} * \mathbf{Meal_{Boolean}} \qquad \text{Eq 1.4.2}$$

$$\mathbf{Protein_{Meal}}[\frac{\mathbf{mg}}{\mathbf{min}}] = \frac{\mathbf{10^3 * Protein_{Amount}[g]}}{\mathbf{Meal_{Length}[min]}} * \mathbf{Meal_{boolean}} \qquad \text{Eq 1.4.3}$$

where $\mathbf{Meal_{boolean}}$ is the previous declared boolean that turns on and off the meal flow, where $\mathbf{MealSolid_{Boolean}}$ and $\mathbf{MealLiquid_{Boolean}}$ indicate the phase of the meal, and where $\mathbf{CarbohydrateAmount}t$ and $\mathbf{Protein_{Amount}}$ are the total amount of respective nutrients in the meal. Equation 1.4.1 – Equation 1.4.3 thus calculate a flow of respective macronutrients during the meal and assume that the meal is consumed at a constant rate.

### Model description: ODEs and reactions

A model reaction corresponds to a chemical reaction or flow of material. The reactions, together with the inputs, events, and parameters, define how the model states change over time. Each ODE describes the time derivative of a specific state (Table 1). All ODEs will henceforth be defined one by one, together with their underlying assumptions. Each ODE is assigned a unique identifier, and all auxiliary equations used in that ODE are assigned identifiers that extend the identifier of the corresponding ODE. For example, the third equation corresponding to ODE numbered X is numbered as X.3.

The first ODE describes the amount of protein in the intestines, and it is a new equation not present in previous models. The state $\mathbf{Intestines_{Protein}}$ describes the pool of protein from a meal in the intestines. The ODE of the state $\mathbf{Intestines_{Protein}}$ is dependent on **Protein**$_{\mathbf{Meal}}$ (Equation 1.4.3), and function $\mathbf{ProteinDigestion}$, which is the rate of digestion from protein into amino acids, see equation 2.

$$\frac{\mathbf{d}}{\mathbf{dt}}(\mathbf{Intestines_{Protein}}) = \left(\frac{\mathbf{Protein_{Meal}}\left[\frac{\mathbf{mg}}{\mathbf{min}}\right]}{\mathbf{BW\ [kg]}}\right) - \left(\mathbf{ProteinDigestion}\ \left[\frac{\mathbf{mg/kg}}{\mathbf{min}}\right]\right) \qquad \text{Eq 2}$$

where the parameter **BW** is the total bodyweight of the simulated person in kg. The function $\mathbf{ProteinDigestion}$ is the product of the parameter $\mathbf{ProteinBreakdown}$ and the amount of protein in the intestines, state $\mathbf{Intestines_{Protein}}$, see Equation 2.1.

$$\mathbf{ProteinDigestion}\left[\frac{\mathbf{mg/kg}}{\mathbf{min}}\right] = \mathbf{ProteinBreakdown}\ \left[\frac{\mathbf{1}}{\mathbf{min}}\right] * \ \mathbf{Intestines_{Protein}}\left[\frac{\mathbf{mg}}{\mathbf{kg}}\right] \qquad \text{Eq 2.1}$$

$\mathbf{ProteinDigestion}$ affects the state $\mathbf{Intestines_{aa}}$ which is the amount of amino acids in the intestines from food intake. The amino acids in the intestines get transported out to the body through the function $\mathbf{aaTransportation_{Meal}}$, see Equation 3.

$$\frac{\mathbf{d}}{\mathbf{dt}}(\mathbf{Intestines_{aa}}) = \mathbf{ProteinDigestion}\ \left[\frac{\mathbf{mg/kg}}{\mathbf{min}}\right] - (\ \mathbf{aaTransportation_{Meal}})\ \left[\frac{\mathbf{mg/kg}}{\mathbf{min}}\right] \qquad \text{Eq 3}$$

The transportation of amino acids from intestines, $\mathbf{aaTransportation_{Meal}}$, is separated into two different flows; i) $\mathbf{aaIntoLiver_{Meal}}$ and ii) $\mathbf{aaIntoBody_{Meal}}$.
The flow $\mathbf{aaIntoLiver_{Meal}}$ is the total flow of amino acids from meals into the liver. The flow $\mathbf{aaIntoBody_{Meal}}$ represents the flow of amino acids from the meal that is not sent into the liver, see equation 3.1.

$$\mathbf{aaTransportation_{Meal}}\left[\frac{\mathbf{mg/kg}}{\mathbf{min}}\right] \ = \ \mathbf{aaIntoLiver_{Meal}}\left[\frac{\mathbf{mg/kg}}{\mathbf{min}}\right] + \ \mathbf{aaIntoBody_{Meal}}\left[\frac{\mathbf{mg/kg}}{\mathbf{min}}\right] \qquad \text{Eq 3.1}$$

The model flow $\mathbf{aaIntoBody_{Meal}}$ is the product of the parameter $\mathbf{ProteinMeal_K}$, which is a general amino acid transportation constant, the total amount of amino acids in the intestines, state $\mathbf{Intestines_{aa}}$, and the function $\mathbf{GlyDep_{MealPositive}}$. $\mathbf{GlyDep_{MealPositive}}$ is in a series of similarly structured functions that uses glycogen in the liver as a global homeostatic regulator. $\mathbf{GlyDep_{MealPositive}}$ upregulates the flow of amino acids out to the body when there is no shortage of stored energy, hence decreasing the relative flow of amino acids to the liver when gluconeogenesis is downregulated. This is further motivated by the increased use of amino acids for anabolic processes outside of the liver when the body is in an anabolic state, see equation 3.1.1 for flow of amino acids to non-liver organs.

$$\mathbf{aaIntoBody_{Meal}}\left[\frac{mg/kg}{min}\right] = \mathbf{ProteinMeal_K}\left[\frac{1}{min}\right] * \mathbf{Intestines_{aa}}\left[\frac{mg}{kg}\right] * \mathbf{GlyDep_{MealPositive}}\left[\frac{mg}{kg}\right] \qquad \text{Eq 3.1.1}$$

The flow $\mathbf{aaIntoBody_{Meal}}$ is upregulated during high energy levels through the function $\mathbf{GlyDep_{MealPositive}}$. This function is defined by the quotient of the amount of hepatic glycogen, state $\mathbf{Gly_L}$, and the parameter $\mathbf{GlyDep_{Meal}}$. The dependency between the flow of amino acids and the energy levels in the liver does not necessarily need to be 1:1, and therefore the parameter $\mathbf{GlyDepEXP_{Meal}}$ is introduced to ensure sufficient flexibility of the model, see Equation 3.1.1.1.

$$\mathbf{GlyDep_{MealPositive}}\left[\frac{mg}{kg}\right] = \left(\frac{\mathbf{Gly_L}\left[\frac{mg}{kg}\right]}{\mathbf{GlyDep_{Meal}}\left[\frac{mg}{kg}\right]}\right)^{\mathbf{GlyDepEXP_{Meal}}} \qquad \text{Eq 3.1.1.1.}$$

As described in equation 3.1, a fraction of the amino acids from ingested food is transported to the liver. The model assumes that these amino acids will either be metabolized to pyruvate or enter the TCA cycle. The amino acid profile of the meal determines which fraction enters each of the two processes and will either be sent into the liver as pyruvate, $\mathbf{aaIntoPyruvate_{Meal}}$, or into the TCA cycle, $\mathbf{aaIntoTCAcycle_{Meal}}$, see equation 3.1.2.

$$\mathbf{aaIntoLiver_{Meal}}\left[\frac{mg/kg}{min}\right] = \mathbf{aaIntoPyruvate_{Meal}}\left[\frac{mg/kg}{min}\right] + \mathbf{aaIntoTCAcycle_{Meal}}\left[\frac{mg/kg}{min}\right] \qquad \text{Eq 3.1.2}$$

The transportation rate of the amino acids into the liver is assumed to be at the same rate as to the rest of the body, hence both flows in Equation 3.1.2 are dependent on the same parameter $\mathbf{ProteinMeal_K}$. The flow of amino acids from the intestines is also dependent on the total amount of amino acids in the intestines, state $\mathbf{Intestines_{aa}}$. The separation of amino acids into the two different flows $\mathbf{aaIntoPyruvate_{Meal}}$ and $\mathbf{aaIntoTCAcycle_{Meal}}$, is controlled by the parameter $\mathbf{Aminoprofile_K}$, see equation 3.1.2.1 and 3.1.2.2.

$$\mathbf{aaIntoPyruvate_{Meal}}\left[\frac{mg/kg}{min}\right] = \mathbf{Aminoprofile_K}\,[\%] * \mathbf{ProteinMeal_K}\left[\frac{1}{min}\right] * \mathbf{Intestines_{aa}}\left[\frac{mg}{kg}\right] * \mathbf{GlyDep_{MealNegative}}\left[\frac{mg}{kg}\right] \qquad \text{Eq 3.1.2.1}$$

$$\mathbf{aaIntoTCAcycle_{Meal}}\left[\frac{mg/kg}{min}\right] = (\mathbf{1} - \mathbf{Aminoprofile_K})\,[\%] * \mathbf{ProteinMeal_K}\left[\frac{1}{min}\right] * \mathbf{Intestines_{aa}}\left[\frac{mg}{kg}\right] * \mathbf{GlyDep_{MealNegative}}\left[\frac{mg}{kg}\right] \qquad \text{Eq 3.1.2.2}$$

where the function $\mathbf{GlyDep_{MealNegative}}$ makes the flow dependent on the fed state of the individual. This is motivated by the upregulation of gluconeogenesis during an unfed state, with an increased need for amino acids in the catabolic processes. The function $\mathbf{GlyDep_{MealNegative}}$ thus introduces a negative dependency on energy levels, and when glycogen is high the flow becomes low. This is modelled using the previously defined parameter $\mathbf{GlyDep_{Meal}}$ (in equation 3.1.1.1) which is divided by the amount of glycogen in the liver, state $\mathbf{Gly_L}$. However, this dependency does not necessarily need to be a ratio of 1:1, and the exponent $\mathbf{GlyDepEXP_{Meal}}$ is introduced to ensure sufficient flexibility of the model. A high value of $\mathbf{GlyDepEXP_{Meal}}$ makes the flow highly dependent on glycogen concentration and vice versa, see Equation 3.1.2.3.

$$\mathbf{GlyDep_{MealNegative}}\left[\frac{mg}{kg}\right] = \left(\frac{\mathbf{GlyDep_{Meal}}\left[\frac{mg}{kg}\right]}{\mathbf{Gly_L}\left[\frac{mg}{kg}\right]}\right)^{\mathbf{GlyDepEXP_{Meal}}} \qquad \text{Eq 3.1.2.3}$$

The next ODE describes the start of the carbohydrate metabolism from a solid meal in the stomach, state **Qsto1,** and is kept intact from the Dalla Man model et al 2007 [1], see Equation 4.

$$\frac{d}{dt}(\mathbf{Qsto1}) = (\mathbf{Carbohydrate_{Solid}})\left[\frac{mg}{min}\right] - (\mathbf{Qsto1toQsto2})\left[\frac{mg}{min}\right] \qquad \text{Eq 4}$$

The function **Qsto1toQsto2** describes the change of the state of ingested carbohydrates. As stated in the original Dalla Man model [1] there is a difference between ingested carbohydrates from food and from a liquid. The carbohydrates from a liquid meal goes directly to the next state, **Qsto2**, and are hence not subjected to the time delay introduced by the ODE of **Qsto1**. The time delay is calculated through multiplying the parameter **Kgri**, with the amount of carbohydrates in the stomach from ingested food. This step is therefore only necessary in the digestion of solid meals, flow $\mathbf{Carbohydrate_{Solid}}$, and not liquids. The function **Qsto1toQsto2** represents the rate of change from this metabolic state and depends on the amount of carbohydrates in the state **Qsto1**, see Equation 4.1.

$$\mathbf{Qsto1ToQsto2}\left[\frac{mg}{min}\right] = \mathbf{K_{gri}}\left[\frac{1}{min}\right] * \mathbf{Qsto1}\ [\mathbf{mg}] \qquad \text{Eq 4.1}$$

The carbohydrates of the flow **Qsto1toQsto2** are further digested in the state **Qsto2**. Carbohydrates that are not going through the extra digestion step is sent into the ODE of the state **Qsto2** through the flow $\mathbf{Carbohydate_{Liquid}}$ which is declared in the event segment, see equation 1.4.2. **Qsto2ToQgut** is a function describing the flow of carbohydrates out of the state **Qsto2**, see Equation 5.

$$\frac{d}{dt}(\mathbf{Qsto2}) = \left(\mathbf{Carbohydrate_{Liquid}} + \mathbf{Qsto1toQsto2}\right)\left[\frac{mg}{min}\right] - (\mathbf{Qsto2ToQgut})\left[\frac{mg}{min}\right] \qquad \text{Eq 5}$$

The carbohydrates in the state **Qsto2** are digested and transported from the state with the function **Qsto2ToQgut**, see Equation 5.1.

$$\mathbf{Qsto2ToQgut}\left[\frac{mg}{min}\right] = \mathbf{K_{empt}}\left[\frac{1}{min}\right] * \mathbf{Qsto2}\left[\frac{1}{mg}\right] \qquad \text{Eq 5.1}$$

The flow of carbohydrates from the stomach, **Qsto2ToQgut**, is partly depending on the function **Kempt,** which is a kept intact from the previous model and describes the gastric emptying of the stomach into the gut, and the amount of carbohydrates in state **Qsto2**. In the function **Kempt**, the flow of carbohydrates is dependent on the size of the ingested meal, parameter **D**, that is declared through an event at the start of the meal, and initially assigned the value of $\mathbf{Meal_{Amount}}$, which is the total size of carbohydrates in mg. The function value of $\mathbf{K_{empt}}$ is dynamic and decreases over time defined by the function **aa**, equation 5.1.2, to the minimum rate of $\mathbf{K_{Min}}$, followed by an increase to $\mathbf{K_{Max}}$ with the rate of the function **cc**, equation 5.1.3. **Qsto** is the sum of carbohydrates in the stomach, equation 5.1.4.
The parameter **b** declares the decreasing rate of change at $\frac{K_{max}-K_{min}}{2}$, and vice versa for the parameter **d** for the increasing rate of change to $\mathbf{K_{Max}}$, see Equation 5.1.1 for function **Kempt**.

$$\mathbf{K_{empt}}\left[\frac{1}{min}\right] = \mathbf{K_{min}} + \frac{\mathbf{K_{max}}-\mathbf{K_{min}}}{2} * \mathbf{tanh}\big(\mathbf{aa} * (\mathbf{Q_{sto}} - \mathbf{b} * \mathbf{D})\big) - \mathbf{tanh}\big(\mathbf{cc} * (\mathbf{Q_{sto}} - \mathbf{d} * \mathbf{D})\big) + \mathbf{2}) \qquad \text{Eq 5.1.1}$$

The transition between the two digestion rates, $\mathbf{K_{Max}}$ and $\mathbf{K_{Min}}$**,** is calculated through the functions **aa** and **cc**. The transitions are dependent on the parameter **D**, total size of the meal, and the parameters **b** and **d**, which are the same parameters as in equation 5.1.1, see Equations 5.1.2 and 5.1.3.

$$\mathbf{aa} = \frac{\mathbf{2.5*D}}{\mathbf{1-b}} \qquad \text{Eq 5.1.2}$$

$$\mathbf{cc} = \frac{\mathbf{2.5*D}}{\mathbf{d}} \qquad \text{Eq 5.1.3}$$

In the calculation of $K_{empt}$, equation 5.1.1, a dependency on $Q_{sto}$ is declared. $Q_{sto}$ is simply the sum of the amount of carbohydrates in the stomach, states $Q_{sto1}$ and $Q_{sto2}$, see equation 5.1.4.

$$\mathbf{Q_{sto}[mg] = Q_{sto1}[mg] + Q_{sto2}[mg]} \qquad \text{Eq 5.1.4}$$

The flow of carbohydrates from the stomach to the gut, flow **Qsto2ToQgut**, is sent into the state **Qgut**, which represents the carbohydrates in the gut from a meal intake. The rate of the metabolism of carbohydrates to glucose is calculated through the function $\mathbf{Qgut_{out}}$. The ODE of state **Qgut** is kept unchanged from previous model, see Equation 6.

$$\frac{d}{dt}(\mathbf{Qgut}) = (\mathbf{Qsto2ToQgut})\left[\frac{mg}{min}\right] - (\mathbf{Qgut_{out}})\left[\frac{mg}{min}\right] \qquad \text{Eq 6}$$

Carbohydrates in state **Qgut** leaves the gut by the flow $\mathbf{Qgut_{Out}}$, defined as the product of the parameter $\mathbf{K_{abs}}$ and the amount of carbohydrates in the gut, state **Qgut**, see Equation 6.1.

$$\mathbf{Qgut_{out}}\left[\frac{mg}{min}\right] = \mathbf{K_{abs}}\left[\frac{1}{min}\right] * \mathbf{Qgut}\,[mg] \qquad \text{Eq 6.1}$$

A substantial portion of the ingested carbohydrates are converted into glucose and transported into the plasma described as state **Gp**. This rate of appearance of glucose from the meal is defined by the flow **Ra**. From the state **Gp**, the glucose uptake of tissue is represented by the flow **GpToGt**. The insulin independent utilization of glucose, for example by the brain, is described by the parameter $\mathbf{U_{ii}}$. Glucose uptake of plasma from tissue is defined by the flow **GtToGp.** This ODE is kept intact from previous model, see Equation 7.

$$\frac{d}{dt}(\mathbf{Gp}) = (\mathbf{Ra} + \mathbf{GtToGp})\left[\frac{mg/kg}{min}\right] - (\mathbf{U_{ii}} + \mathbf{GpToGt})\left[\frac{mg/kg}{min}\right] \qquad \text{Eq 7}$$

In the function **Ra** the parameter **f** is introduced to account for losses from the ingested carbohydrates and the rate of appearance in plasma. The parameter **f** is defined as a percentage, and it is multiplied with the flow out of the state **Qgut** to get **Ra**. In addition, the function **Ra** is divided by the total body weight, **BW** to get the right unit , see Equation 7.1.

$$\mathbf{Ra}\left[\frac{mg/kg}{min}\right] = \mathbf{f}[\%] * \frac{\mathbf{Qgut_{out}}\left[\frac{mg}{min}\right]}{\mathbf{BW}\,[kg]} \qquad \text{Eq 7.1}$$

The flow of glucose from the plasma into tissue is calculated as the product of the parameter $\mathbf{K_1}$ and the amount of glucose in plasma state **Gp**, see Equation 7.2.

$$\mathbf{GpToGt}\left[\frac{mg/kg}{min}\right] = \mathbf{K_1}\left[\frac{1}{min}\right] * \mathbf{Gp}\left[\frac{mg}{kg}\right] \qquad \text{Eq 7.2}$$

The glucose uptake from the tissue into the plasma is the product of the parameter $\mathbf{K_2}$ and the total glucose in tissue, state **Gt**, see Equation 7.3.

$$\mathbf{GtToGp}\left[\frac{mg/kg}{min}\right] = \mathbf{K_2}\left[\frac{1}{min}\right] * \mathbf{Gt}\left[\frac{mg}{kg}\right] \qquad \text{Eq 7.3}$$

The insulin independent utilization of glucose is modelled by the parameter $\mathbf{U_{ii}}$, which represents the constant need of glucose from cells that cannot use other fuel sources, such as ketone bodies, even during longer periods of fasting.

All glucose in tissue is summarized as the state **Gt.** Glucose is transported into tissue from plasma and by the endogenous glucose production, summarized as the flow **EGP**. Compared to the previous model of **EGP**, our model has been extended in the following ways: the physical interpretation has been changed from a non-physiological flow of glucose from an inexhaustible source into the model in form of a constant value minus the current external source of glucose, to a physical diffusion-driven flow directly influenced by glucose production from glycogenolysis and gluconeogenesis. **EGP** is also expanded to include glucose production both from the liver and the kidneys and the total **EGP** is the sum of glucose produced from both organs. An insulin-dependent utilization of glucose in tissue is represented by the function $\mathbf{U_{id}}$. The transport of glucose into the liver is defined as $\mathbf{U_{iil}}$, insulin independent utilization of liver, see the ODE of the state **Gt** in Equation 8.

$$\frac{\mathbf{d}}{\mathbf{dt}}(\mathbf{Gt}) = (\ \mathbf{GpToGt} + \mathbf{EGP}\ )\left[\frac{\mathbf{mg/kg}}{\mathbf{min}}\right] - (\ \mathbf{U_{id}} +\ \mathbf{GtToGp} +\ \mathbf{U_{iil}})\ \left[\frac{\mathbf{mg/kg}}{\mathbf{min}}\right] \qquad \text{Eq 8}$$

The endogenous glucose production, function **EGP**, is the sum of the production from the liver $\mathbf{EGP_{Liver}}$ and the kidneys $\mathbf{EGP_{Kidneys}}$, see equation 8.1.

$$\mathbf{EGP}\left[\frac{\mathbf{mg/kg}}{\mathbf{min}}\right] = \mathbf{EGP_{Liver}}\left[\frac{\mathbf{mg/kg}}{\mathbf{min}}\right] + \mathbf{EGP_{Kidneys}}\left[\frac{\mathbf{mg/kg}}{\mathbf{min}}\right] \qquad \text{Eq 8.1}$$

The glucose production in the kidneys, function $\mathbf{EGP_{Kidneys}}$, is dependent on the fed state of the individual via the function $\mathbf{GlyDep_{inflow}}$. This factor is multiplied by the parameter $\mathbf{EGP_{KidneysK}}$ to determine the magnitude of the gluconeogenesis in the kidneys, see Equation 8.1.1.

$$\mathbf{EGP_{Kidneys}}\left[\frac{\mathbf{mg/kg}}{\mathbf{min}}\right] = \mathbf{EGP_{KidneysK}}\left[\frac{\mathbf{1}}{\mathbf{min}}\right] * \mathbf{GlyDep_{inflow}}\left[\frac{\mathbf{mg}}{\mathbf{kg}}\right] \qquad \text{Eq 8.1.1}$$

The gluconeogenesis is upregulated during time periods of low energy levels. This regulation is driven by the function $\mathbf{GlyDep_{inflow}}$ defined by the quotient between the parameter $\mathbf{GlyDepIn_K}$ and$\mathbf{Gly_L}$. As previously motivated for similar homeostatic regulations of hepatic glycogen, this dependency does not need to be a ratio of 1:1, hence the exponent $\mathbf{GlyDepIn_{kEGP}}$ is added to ensure model flexibility, see Equation 8.1.1.1.

$$\mathbf{GlyDep_{inflow}}\left[\frac{\mathbf{mg}}{\mathbf{kg}}\right] = \left(\frac{\mathbf{GlyDepIn_K}\left[\frac{\mathbf{mg}}{\mathbf{kg}}\right]}{\mathbf{Gly_L}\left[\frac{\mathbf{mg}}{\mathbf{kg}}\right]}\right)^{\mathbf{GlyDepIn_{kEGP}}} \qquad \text{Eq 8.1.1.1.}$$

In the equations of **EGP**, Equation 8.1, the glucose production from the liver is defined as $\mathbf{EGP_{Liver}}$. Glucose is transported into and out of the liver by the transport protein GLUT2. The amount of GLUT2 in the liver is limited, and the flow out of the liver, $\mathbf{EGP_{Liver}}$, and into the liver, $\mathbf{U_{iil}}$, can therefore be saturated. This saturation is modelled by the maximum flow parameters $\mathbf{EGPLiver_{MAX}}$ and $\mathbf{Uiil_{MAX}}$. The rate is defined by the parameter $\mathbf{EGPLiver_0}$. The diffusion rate of glucose from the liver into tissue is also dependent on the amount of glucose in the liver, state **Gl**, see Equation 8.1.2.

$$\mathbf{EGP_{Liver}}\left[\frac{\mathbf{mg/kg}}{\mathbf{min}}\right] = \frac{\mathbf{EGPLiver_{MAX}}\left[\frac{\mathbf{1}}{\mathbf{min}}\right] * \mathbf{Gl}\left[\frac{\mathbf{mg}}{\mathbf{kg}}\right]}{\mathbf{EGPLiver_0}\left[\frac{\mathbf{1}}{\mathbf{min}}\right] + \mathbf{Gl}\left[\frac{\mathbf{mg}}{\mathbf{kg}}\right]} \qquad \text{Eq 8.1.2}$$

The diffusion rate of glucose from tissue into liver is dependent on the amount of glucose in tissue, $\mathbf{Gt}$ and the same glucose transporter, GLUT2, as the transport of glucose out of the liver. To take into consideration that the state **Gt** is the sum of all glucose in tissue, different sets of parameters must be used to describe the two directions of GLUT2 transport. The maximum flow of glucose uptake of the liver is represented by the parameter $\mathbf{Uiil_{MAX}}$ and the maximum rate defined by the parameter $\mathbf{Uiil_0}$, see Equation 8.3

$$\mathbf{U_{iil}}\left[\frac{\mathbf{mg/kg}}{\mathbf{min}}\right] = \frac{\mathbf{Uiil_{MAX}}\left[\frac{\mathbf{1}}{\mathbf{min}}\right] * \mathbf{Gt}\left[\frac{\mathbf{mg}}{\mathbf{kg}}\right]}{\mathbf{Uiil_0}\left[\frac{\mathbf{1}}{\mathbf{min}}\right] + \mathbf{Gt}\left[\frac{\mathbf{mg}}{\mathbf{kg}}\right]} \qquad \text{Eq 8.3}$$

In the ODE describing glucose in tissue, state **Gt** (Equation 8), the insulin dependent utilization is defined by the function $\mathbf{U_{id}}$. This function is composed of a flow of glucose both into muscle tissue, $\mathbf{U_{idm}}$, and into adipocytes, $\mathbf{U_{idf}}$, see Equation 8.3.1.

$$\mathbf{U_{id}}\left[\frac{\mathbf{mg/kg}}{\mathbf{min}}\right] = \mathbf{U_{idf}}\left[\frac{\mathbf{mg/kg}}{\mathbf{min}}\right] + \mathbf{U_{idm}}\left[\frac{\mathbf{mg/kg}}{\mathbf{min}}\right] \qquad \text{Eq 8.3.1}$$

The maximum utilization of glucose in adipocytes is determined by the function $\mathbf{Vf_{MAX}}$ and the maximum utilization of glucose in muscle is determined by the function $\mathbf{Vm_{MAX}}$. Both utilization functions are dependent on the amount of glucose in tissue, state **Gt,** where the half of the maximum velocity, $\mathbf{V_{Max}}$, is reached when the value of the state **Gt** is the same value as $\mathbf{K_{f0}}$ and $\mathbf{K_{m0}}$ respectively**,** see Equation 8.3.1.2 and 8.3.1.2.

$$\mathbf{U_{idf}}\left[\frac{\mathbf{mg/kg}}{\mathbf{min}}\right] = \frac{\mathbf{Vf_{MAX}*Gt}\left[\frac{\mathbf{mg}}{\mathbf{kg}}\right]}{\mathbf{Kf_0+\ Gt}\left[\frac{\mathbf{mg}}{\mathbf{kg}}\right]} \qquad \text{Eq 8.3.1.1}$$

$$\mathbf{U_{idm}}\left[\frac{\mathbf{mg/kg}}{\mathbf{min}}\right] = \frac{\mathbf{Vm_{MAX}*Gt}\left[\frac{\mathbf{mg}}{\mathbf{kg}}\right]}{\mathbf{Km_0+\ Gt}\left[\frac{\mathbf{mg}}{\mathbf{kg}}\right]} \qquad \text{Eq 8.3.1.2}$$

The maximum reaction rates are given by the two functions $\mathbf{Vf_{Max}}$ and $\mathbf{Vm_{Max}}$. These maximum flow functions are dependent on a basal value, $\mathbf{V_{f0}}$ and $\mathbf{V_{m0}}$, and a dynamic correlation to the insulin response in tissue, **InsulinResponseTissue**. This dynamic correlation is calibrated by the parameters $\mathbf{V_{fX}}$ and $\mathbf{V_{mX}}$.

One limitation of previous models is that they do not scale well between small (<500 kcal) and big meals (>500 kcal). The reason for this may be that the model neglects glucose homeostatic systems that may play a bigger role during hyperglycemia, for example renal exclusion, or that the glucose rate of appearance from the intestines is overestimated for big meals that may have a slower rate of appearance. Currently, we lack fundamental data to create models representing such hypotheses at a mechanistic level. To address the scalability problem with the least amount of non-physiological changes to the model, we introduce the parameter $\mathbf{InsulinDep_{exp}}$ to increase the utilization of glucose during big meals by a nonlinear relation between usage of glucose in relation to insulin response in tissue. In future work we propose to solve the scalability problem either by expanding the carbohydrate metabolism, or introduce new systems, for example a dynamic glucose uptake by the brain and kidneys. The function **InsulinResponseTissue** represent the insulin response of the individual or population and may thus be decreased to represent individuals with high insulin resistance, thus enabling simulations of individuals with metabolic syndromes like type 2 diabetes, see Equation 8.3.1.3 and Equation 8.3.1.4

$$\mathbf{V_{fmax}}\left[\frac{\mathbf{1}}{\mathrm{min}}\right] = (\mathbf{V_{f0}}\left[\frac{\mathbf{1}}{\mathrm{min}}\right] + \mathbf{V_{fX}}\left[\frac{\mathbf{1}}{\mathrm{min}}\right] * \mathbf{InsulinResponseTissue})^{\mathbf{InsulinDep_{exp}}} \qquad \text{Eq 8.3.1.3}$$

$$\mathbf{V_{mmax}}\left[\frac{\mathbf{1}}{\mathrm{min}}\right] = (\mathbf{V_{m0}}\left[\frac{\mathbf{1}}{\mathrm{min}}\right] + \mathbf{V_{mX}}\left[\frac{\mathbf{1}}{\mathrm{min}}\right] * \mathbf{InsulinResponseTissue})^{\mathbf{InsulinDep_{exp}}} \qquad \text{Eq 8.3.1.4}$$

A delay between the first appearance of insulin in the blood and the insulin response in tissue is kept unchanged from the previous model [1]. The rate of the delay is determined by the parameter $\mathbf{It_{delayK}}$. A high value of $\mathbf{It_{delayK}}$ results in a fast response, while a low value of the parameter $\mathbf{It_{delayK}}$ decreases the amplitude of the insulin in tissue response and lengthens it. The reaction **InsulinResponseTissue** is a delay of current insulin in the blood which is described with the state **Ip** [pmol/kg]. The state **Ip** is converted into the unit [pmol/L] through the model variable $\mathbf{Insulin_{Blood}}$, declared in equation 15.2, see Equation 8.3.1.5.

$$\frac{\mathbf{d}}{\mathbf{dt}}\mathbf{InsulinResponseTissue} = \mathbf{It_{delayK}} * (\mathbf{Insulin_{Blood}} - \mathbf{InsulinResponseTissue}) \qquad \text{Eq 8.3.1.5}$$

As previously stated, the amount of glucose in tissue is closely linked to glucose in the liver through the flows of glucose into the liver, $\mathbf{U_{iil}}$, and the flow of glucose from the liver, $\mathbf{EGP_{Liver}}$. The amount of glucose in the liver is represented by the state **Gl**.

In the ODE of the state **Gl,** synthesis of glycogen is represented as the model reaction **Glycogenesis** and the breakdown of glycogen into glucose by the model reaction **Glycogenolysis**. Glucose may be converted into pyruvate through the model reaction **Glycolysis** and synthesized from the TCA cycle by the catabolic reaction **Gluconeogenesis**, see Equation 9 for ODE of state **Gl**.

$$\frac{d}{dt}(\mathbf{Gl}) = (\mathbf{U_{iil}} + \mathbf{Glycogenolysis} + \mathbf{Gluconeogenesis})\left[\frac{mg/kg}{min}\right] - (\mathbf{Glycogenesis} + \mathbf{EGP_{Liver}} + \mathbf{Glycolysis})\left[\frac{mg/kg}{min}\right] \qquad \text{Eq 9}$$

Excess energy can be stored in the liver in the form of glycogen. This process, **Glycogenesis**, is regulated through the anabolic hormone insulin. In the model, **Glycogenesis** is therefore dependent on the insulin response in the liver, represented by the function $\mathbf{InsulinResponse_{Liver}}$.

The maximum reaction rate of **Glycogenesis** is set by the product of the parameter $\mathbf{Vglys_{Max}}$ and the function **InsulinResponse_L.** The model reaction **Glycogenesis** also depends on the amount of glucose in the liver, state **Gl**, where the half of the maximum velocity, is reached when the value of the state **Gl** equals the value of the parameter $\mathbf{VglyS_0}$**,** see equation 9.1.

$$\mathbf{Glycogenesis}\left[\frac{mg/kg}{min}\right] = \frac{\mathbf{VglyS_{max}}\left[\frac{1}{min}\right] * \mathbf{InsulinResponse_L} * \mathbf{Gl}\left[\frac{mg}{kg}\right]}{\mathbf{VglyS_0}\left[\frac{1}{min}\right] + \mathbf{Gl}\left[\frac{mg}{kg}\right]} \qquad \text{Eq 9.1}$$

**Glycogenesis** is regulated by insulin and depends on the function $\mathbf{InsulinResponse_L}$. In the original Dalla man model, a state called **Il** represents hepatic insulin and was used as an intermediate step between insulin in the portal vein and the blood plasma. In our model we have implemented this to have a physiological meaning with metabolic fluxes directly being dependent on the insulin in the liver. This insulin response therefore both affects the synthesis of glycogen and amino acids.

The function $\mathbf{InsulinResponse_L}$ represents the magnitude of the insulin response by the product of the insulin in the liver, the state **Il**, and the parameter$\mathbf{InsulinLiverResponse_K}$￼, see Equation 9.1.1.

$$\mathbf{InsulinResponse_L} = \mathbf{Il}\left[\frac{pmol}{kg}\right] * \mathbf{InsulinLiverResponse_K} \qquad \text{Eq 9.1.1}$$

When there is no external supply of glucose, the body uses the stored energy source glycogen, and converts it into glucose through the reaction of **Glycogenolysis**. The maximum rate of the reaction **Glycogenolysis** is described by the parameter $\mathbf{VglyB_{Max}}$. The model reaction **Glycogenolysis** is dependent on the amount of glucose in the liver, state **Gl,** where half of the maximum velocity is reached when the value of the state **Gl** equals the parameter $\mathbf{VglyB_0}$**,** see Equation 9.2.

$$\mathbf{Glycogenolysis}\left[\frac{mg/kg}{min}\right] = \frac{\mathbf{VglyB_{max}}\left[\frac{1}{min}\right] * \mathbf{Gl}\left[\frac{mg}{kg}\right]}{\mathbf{VglyB_0}\left[\frac{1}{min}\right] + \mathbf{Gl}\left[\frac{mg}{kg}\right]} \qquad \text{Eq 9.2}$$

In the liver, glucose is also converted into pyruvate through the anabolic process called **Glycolysis**. This reaction is determined by the product of the parameter $\mathbf{Glycolysis_k}$**,** the amount of glucose in liver, state **Gl**, and the function $\mathbf{Insulin\ Response_{Liver}}$. The function $\mathbf{InsulinResponse_{Liver}}$ makes the metabolic reaction **Glycolysis** dependent on insulin. $\mathbf{InsulinResponse_{Liver}}$ is the same function used to activate other anabolic reactions in the liver like **Glycogenesis**. To allow the metabolic reactions to have different dependency on insulin, the reaction-specific exponent $\mathbf{GlycolysisEXP_K}$ is added. This is one of many parameters that can be set to calibrate the model to individuals with metabolic syndrome that alters the insulin responsiveness, see Equation 9.3.

$$\mathbf{Glycolysis}\left[\frac{\mathbf{mg/kg}}{\mathbf{min}}\right] = \mathbf{Glycolysis_k}\left[\frac{\mathbf{1}}{\mathbf{min}}\right] * \mathbf{InsulinResponse_{Liver}^{GlycolysisEXP_K}} * \mathbf{Gl}\left[\frac{\mathbf{mg}}{\mathbf{kg}}\right] \quad \text{Eq 9.3}$$

$\mathbf{InsulinResponse_{Liver}}$ has previously been declared in Equation 9.1.1.

When glycogen stores are depleted the insulin-dependent utilization of glucose is lowered and the process of breaking down protein to glucose, **Gluconeogenesis**, is upregulated. **Gluconeogenesis** represents the catabolic reaction of synthesizing glucose from amino acids in the TCA cycle. In the TCA cycle, the component oxaloacetate can be metabolized into phosphoenolpyruvic acid (PEP) and finally be metabolized into glucose. In the model, **Gluconeogenesis** in the liver is hence represented by a flow between a state describing the TCA cycle, state **TCAcycle**$_{\mathbf{L}}$, into glucose in the liver, state **Gl**, see Equation 9.4.

$$\mathbf{Gluconeogenesis}\left[\frac{\mathbf{mg/kg}}{\mathbf{min}}\right] = \mathbf{TCAcycle_L}\left[\frac{\mathbf{mg}}{\mathbf{kg}}\right] * \mathbf{Gluconeogenesis_{TCAk}}\left[\frac{\mathbf{1}}{\mathbf{min}}\right] * \mathbf{GlyDep_{GluconeogenesisNegative}}\left[\frac{\mathbf{mg}}{\mathbf{kg}}\right] \quad \text{Eq 9.4}$$

The model reaction $\mathbf{Gluconeogenesis}$ depends on the previously declared global homeostatic regulator hepatic glycogen, state **GlyL**. This dependency is implemented through the multiplication with the function $\mathbf{GlyDep_{GluconeogenesisNegative}}$, defined in Equation 9.4.1.

$$\mathbf{GlyDep_{GluconeogenesisNegative}} = \left(\frac{\mathbf{GlyDep_{Gluconeogenesis}}\left[\frac{\mathbf{mg}}{\mathbf{kg}}\right]}{\mathbf{Gly_L}\left[\frac{\mathbf{mg}}{\mathbf{kg}}\right]}\right)^{\mathbf{GlyDepEXP_{Gluconeogenesis}}} \quad \text{Eq 9.4.1.}$$

The liver is responsible for a large portion of the synthesis of amino acids in the body. The process of converting glucose into pyruvate, **Glycolysis** (Equation 9.3), is hence sent into the state $\mathbf{Pyruvate_L}$, that represents the amount of pyruvate in the body. Pyruvate in the liver is produced either from **Glycolysis**, a flow from the body called $\mathbf{PyruvateTranslocase}$, or from a meal, flow $\mathbf{aaIntoPyruvate_{Meal}}$. Pyruvate can be oxidized to AcetylCoA through the function $\mathbf{PyruvateOxi_L}$ and be metabolized into glucose through the function $\mathbf{Gluconeogenesis_{Pyruvate}}$, see Equation 10 for ODE of the model state $\mathbf{Pyruvate_L}$.

$$\frac{\mathbf{d}}{\mathbf{dt}}(\mathbf{Pyruvate_L}) = (\mathbf{aaIntoPyruvate_{Meal}} + \mathbf{Glycolysis} + \mathbf{PyruvateTranslocase}) - (\mathbf{PyruvateOxi_L})\left[\frac{\mathbf{mg/kg}}{\mathbf{min}}\right] \quad \text{Eq 10.}$$

The flow of pyruvate from the body into the liver is described through the function $\mathbf{PyruvateTranslocase}$. The function $\mathbf{PyruvateTranslocase}$ is the product of the parameter $\mathbf{PyruvateTranslocase_K}$ and a dynamic relation to our global homeostatic regulator, function $\mathbf{GlyDep_{inflow}}$. This relation to glycogen is set to the same relation as the other flow into the model used in the calculation of glucose from kidneys. This was done to reduce the amount of model parameters, so that the model does not get more complicated then needed. The previous declared function $\mathbf{GlyDep_{inflow}}$ (equation 8.1.1.1) is thus reused, see Equation 10.1.

$$\mathbf{PyruvateTranslocase} = \mathbf{PyruvateTranslocase_K} * \mathbf{GlyDep_{inflow}} \quad \text{Eq 10.1}$$

The oxidation of pyruvate into AcetylCoA, $\mathbf{PyruvateOxi_L}$, is modelled by the product of the parameter $\mathbf{PyruvateOxi_K}$ and the total amount of pyruvate in the liver, state $\mathbf{Pyruvate_L}$. This means that the oxidation is directly proportional to the amount of pyruvate in liver with the magnitude of the parameter $\mathbf{PyruvateOxi_K}$. This is a simplification of the true biology, but a more complex relation would be hard to justify without access to more data on pyruvate and AcetylCoA. The relationship of this oxidation on pyruvate into AcetylCoA is given in Equation 10.2.

$$\mathbf{PyruvateOxi_L}\left[\frac{\mathbf{mg/kg}}{\mathbf{min}}\right] = \mathbf{Pyruvate_L}\left[\frac{\mathbf{1}}{\mathbf{min}}\right] * \mathbf{PyruvateOxi_K}\left[\frac{\mathbf{mg}}{\mathbf{kg}}\right] \qquad \text{Eq 10.2.}$$

The model rection $\mathbf{PyruvateOxi_L}$ describes the oxidation of pyruvate into AcetylCoa. AcetylCoA is a component of the TCA cycle and to avoid unnecessary model complexity the TCA cycle is represented by a single state, $\mathbf{TCAcycle_L}$. This simplification may impact the model behaviour where now all components of the cycle behave the same and amino acids that enter early in the cycle will either be utilized, function $\mathbf{TCAusage}$, or metabolized into glucose, $\mathbf{Gluconeogenesis_{TCAcycle}}$, with the same rate and timing as amino acids enter the cycle in a later phase. We note that this model part must be expanded to represent alternative fuel sources such as ketones and fatty acids. However, for the use of simulating glucose/insulin/glycogen dynamics the simple representation is useful.

In the ODE of the state $\mathbf{TCAcycle_L}$ components enter either from a meal, $\mathbf{aaIntoTCAcycle_{Meal}}$, or from pyruvate $\mathbf{PyruvateOxi_L}$, see Equation 11 for ODE of $\mathbf{TCAcycle_L}$.

$$\frac{\mathbf{d}}{\mathbf{dt}}(\mathbf{TCAcycle_L}) = (\mathbf{aaIntoTCAcycle_{Meal}} + \mathbf{PyruvateOxi_L})\left[\frac{\mathbf{mg/kg}}{\mathbf{min}}\right] - \left(\mathbf{Gluconeogenesis} + \mathbf{TCA_{usage}}\right)\left[\frac{\mathbf{mg/kg}}{\mathbf{min}}\right] \qquad \text{Eq 11.}$$

The utilization of components of the TCA cycle is defined by the function $\mathbf{TCA_{usage}}$. The function $\mathbf{TCA_{usage}}$ is a product of the parameter $\mathbf{TCAusage_K}$, the amount of components in the TCA cycle, state $\mathbf{TCAcycle_L}$ and function $\mathbf{GlyDep_{UtilizationPositive}}$, see Equation 11.1.

$$\mathbf{TCA_{usage}}\left[\frac{\frac{\mathbf{mg}}{\mathbf{kg}}}{\mathbf{min}}\right] = \mathbf{TCAcycle_L}\left[\frac{\mathbf{mg}}{\mathbf{kg}}\right] * \mathbf{TCAusage_K}\left[\frac{\mathbf{1}}{\mathbf{min}}\right] * \mathbf{GlyDep_{UtilizationPositive}} \qquad \text{Eq 11.1.}$$

The function $\mathbf{GlyDep_{UtilizationPositive}}$ reduces wasted energy during catabolism and is constructed like all other functions regulated by the global energy homeostatic regulator glycogen. The physiological reason for this function is to preserve energy when glycogen stores in the liver are low. When this is the case, the body prioritize vital processes for survival, *i.e.* energy for the brain, that cannot use ketone bodies as fuel, and slowly deprioritizes the lesser important functions, such as muscle synthesis. To represent this dependency in the model, the hepatic glycogen, state **Gl**, is divided by the parameter $\mathbf{GlyDep_{Utilization}}$ and the dependency to glycogen is calibrated through the parameter$\mathbf{GlyDepEXP_{Utilization}}$, see equation 11.1.1.

$$\mathbf{GlyDep_{UtilizationPositive}}\left[\frac{\mathbf{mg}}{\mathbf{kg}}\right] = \left(\frac{\mathbf{Gl}\left[\frac{\mathbf{mg}}{\mathbf{kg}}\right]}{\mathbf{GlyDep_{Utilization}}\left[\frac{\mathbf{mg}}{\mathbf{kg}}\right]}\right)^{\mathbf{GlyDepEXP_{Utilization}}} \qquad \text{Eq 11.1.1}$$

All other reactions in the ODE of state $\mathbf{TCAcycle_L}$ (Equation 11), has previously been defined; $\mathbf{aaIntoTCAcycle_{Meal}}$ (Equation 3.1.2.2) and $\mathbf{PyruvateOxi_L}$ (Equation 10.1).

When the body has excess glucose in the blood, hyperglycemia, the production of insulin in the pancreas is upregulated and is represented in model as function **Spo**. From the pancreas the insulin is transported into the portal vein. Insulin concentratioin in the portal vein is is represented by the state **Ipo**. From the portal vein insulin is secreted through the function **S**, see Equation 12 for the ODE of the state **Ipo**.

$$\frac{\mathbf{d}}{\mathbf{dt}}(\mathbf{Ipo}) = \mathbf{Spo}\left[\frac{\mathbf{pmol/kg}}{\mathbf{min}}\right] - \mathbf{S}\left[\frac{\mathbf{pmol/kg}}{\mathbf{min}}\right] \qquad \text{Eq 12.}$$

The model function **Spo** describes the rate of appearance of insulin in the portal vein. **Spo** depends on three different parts: i) Rate of change of glucose in plasma, which is function **ChangeInGlucose** ii) glucose relation to a basal value which is **InsulinStabilization**, and iii) a basal insulin production, which is parameter $\mathbf{S_b}$, see Equation 12.1.

$$\mathbf{S_{po}}\left[\frac{\mathbf{pmol/kg}}{\mathbf{min}}\right] = \mathbf{ChangeInGlucose}\left[\frac{\mathbf{pmol/kg}}{\mathbf{min}}\right] + \mathbf{InsulinStabilization}\left[\frac{\mathbf{pmol/kg}}{\mathbf{min}}\right] + \mathbf{S_b}\left[\frac{\mathbf{pmol/kg}}{\mathbf{min}}\right] \qquad \text{Eq 12.1}$$

The function **InsulinStabilization** represents production based on the difference between glucose in plasma, state **Gp**, and the parameter $\mathbf{G_b}$, which is the basal value of glucose between meals. The function **InsulinStabilization** is implemented as an ODE which causes a delay between an increase in glucose and the insulin production response. The size of the insulin response is determined by the parameter **beta**, and the delay is determined by the parameter **alpha**. Lowering and increasing the parameter **beta** can henceforth be a part of calibrating insulin production response between individuals, enabling simulations of people with metabolic syndromes such as type 2 diabetes. The glucose in plasma is represented by the state **Gp** and because the unit of **Gp** is [mg/kg], the function **Spo** is divided by the molecular weight of glucose to change unit to [mmol/kg]. The parameter **beta** determines the amplitude of the response, and the unit conversion from 'milli' to 'pico' is implicitly part of the parameter value. The function **InsulinStabilization** tries to balance the glucose level to the setpoint **Gb** between meals, see Equation 12.1.1.

$$\frac{\mathbf{d}}{\mathbf{dt}}\mathbf{InsulinStabilization}\left[\frac{\mathbf{pmol/kg}}{\mathbf{min}}\right] = \frac{\mathbf{alpha}}{\mathbf{180.16}} * (\mathbf{beta} * (\mathbf{Gp} - \mathbf{G_b}) - \mathbf{InsulinStabilization}) \qquad \text{Eq 12.1.1}$$

Balancing glucose is of high importance for the body and regulation of energy. Too high glucose levels increase the risk of haemoglobin, Hb, spontaneously binding to glucose creating HbA1C. HbA1C binds to glucose with high affinity and makes hemoglobin unable to transport oxygen over its lifetime. A high stabilization force, meaning a high value of the parameter **beta**, results in an effective control of both insulin and glucose. Naturally this parameter is expected to be lower in a diabetic patient.

The second part of the **Spo** calculations is the function **ChangeInGlucose**, which calculates the insulin production based on change of plasma glucose, the time derivative of the state **Gp**. Model reaction **GpToGt** (Equation 7.2), and parameter **Uii**, which is the insulin independent utilization of glucose, is the sum of all negative rates of change. Model flows **Ra** (Equation 7.1), and **GtToGp** (Equation 7.3) is the positive rate of changes of glucose. The amplitude of the insulin production response that is dependent to this previously mentioned rate of change is ultimately scaled by the parameter **K**, which is divided by the molecular weight of glucose to change unit from [mg/kg] to [mmol/kg], see Equation 12.1.2.

$$\mathbf{Change\ in\ Glucose}\left[\frac{\mathbf{mg/kg}}{\mathbf{min}}\right] = \frac{\mathbf{K}}{\mathbf{180.16}} * (\mathbf{Ra} + \mathbf{GtToGp} - \mathbf{U_{ii}} - \mathbf{GpToGt})\left[\frac{\mathbf{mg/kg}}{\mathbf{min}}\right] \qquad \text{Eq 12.1.2.}$$

As previously mentioned, the third and final part of the insulin production is based on a basal insulin production that is determined by the parameter $\mathbf{S_b}$.

The rate of disappearance of insulin from the portal vein, **S** (Equation 12), depends on the secretion of insulin from portal vein into the liver. This secretion is defined by the product of the parameter **gamma** and the total insulin in the portal vein, state **Ipo**, see Equation 12.2.

$$\mathbf{S}\left[\frac{\mathbf{pmol/kg}}{\mathbf{min}}\right] = \mathbf{gamma}\left[\frac{\mathbf{1}}{\mathbf{min}}\right] * \mathbf{Ipo}\left[\frac{\mathbf{pmol}}{\mathbf{kg}}\right] \qquad \text{Eq 12.2}$$

The flow **S** goes from the state **Ipo**, insulin in portal vein, to the state **Il**, insulin in liver. The ODE of **Il** was present in the previous model but did not affect any metabolic reactions in the model and simply worked as a delay between insulin production and the observed insulin concentration in circulation. The ODE of state **Il** is kept intact from the previous model, but the parameter values have been revised to reflect that the state **Il** now affects hepatic metabolic reactions such as synthesis of both glycogen and pyruvate. The flow **IpToIl** describes the diffusion of insulin from the blood plasma to the liver and the flow $\mathbf{IlToIp}$ describes the flow back into the blood. $\mathbf{InsulinDegradation_{Liver}}$ describes insulin degradation in the liver, see Equation 13 for ODE of state **Il**.

$$\frac{d}{dt}(\mathbf{Il}) = (\boldsymbol{S} + \mathbf{IpToIl})\left[\frac{\mathbf{pmol/kg}}{\mathbf{min}}\right] - (\mathbf{IlToIp} + \mathbf{InsulinDegradation_{Liver}})\left[\frac{\mathbf{pmol/kg}}{\mathbf{min}}\right] \quad \text{Eq 13}$$

The transport of insulin between the liver and the blood, $\mathbf{LiverToBlood_{Insulin}}$, is defined by the product of the parameter $\mathbf{m_1}$ and the total current amount of insulin in the liver, state **Il**, see Equation 13.1.

$$\mathbf{IpToIl}\left[\frac{\mathbf{pmol/kg}}{\mathbf{min}}\right] = \mathbf{m_1}\left[\frac{\mathbf{1}}{\mathbf{min}}\right] * \mathbf{Il}\left[\frac{\mathbf{pmol}}{\mathbf{kg}}\right] \quad \text{Eq 13.1}$$

The transportation from the blood plasma to the liver is described by the flow $\mathbf{IlToIp}$. The function $\mathbf{IlToIp}$ is defined by the product of the parameter $\mathbf{m_2}$ and the amount of insulin in plasma, state **Ip**, see Equation 13.2.

$$\mathbf{IlToIp}\left[\frac{\mathbf{pmol/kg}}{\mathbf{min}}\right] = \mathbf{m_2}\left[\frac{\mathbf{1}}{\mathbf{min}}\right] * \mathbf{Ip}\left[\frac{\mathbf{pmol}}{\mathbf{kg}}\right] \quad \text{Eq 13.2}$$

The liver is the main organ for insulin degradation and approximately 50% of insulin from the portal vein never leaves the liver. This degradation is modelled by the function $\mathbf{InsulinDegradation_{Liver}}$ defined by the product of the model function $\mathbf{M_3}$ and the insulin in the liver, state **Il**. Function $\mathbf{InsulinDegradation_{Liver}}$ and the subfunctions, $\mathbf{M_3}$ and **HepaticExtraction** are kept intact from the previous model, see Equation 13.3.

$$\mathbf{InsulinDegradation_{Liver}}\left[\frac{\mathbf{pmol/kg}}{\mathbf{min}}\right] = \mathbf{M_3}\left[\frac{\mathbf{1}}{\mathbf{min}}\right] * \mathbf{Insulin_{Liver}}\left[\frac{\mathbf{pmol}}{\mathbf{kg}}\right] \quad \text{Eq 13.3}$$

The insulin degradation in the liver, function $\mathbf{M_3}$, depends on the parameter $\mathbf{m_1}$ and the function **HepaticExtraction**, see Equation 13.3.1.

$$\mathbf{M_3}\left[\frac{\mathbf{1}}{\mathbf{min}}\right] = \frac{\mathbf{HepaticExtraction} * \mathbf{m_1}}{\mathbf{1 - HepaticExtraction}} \quad \text{Eq 13.3.1}$$

The model function **HepaticExtraction** decreases the degradation during increased secretion of insulin into the liver. This results in a short delay between the increased rate of change of insulin into the liver and the insulin degradation. **HepaticExtraction** is separated into a dependency on secretion of insulin, **S**, and a basal value determined by the parameter $\mathbf{m_6}$. The dependency on the secretion of insulin, **S**, is determined by the parameter $\mathbf{m_5}$, see Equation 13.3.2.

$$\mathbf{HepaticExtraction} = -\mathbf{m_5} * \mathbf{S} + \mathbf{m_6} \quad \text{Eq 13.3.2}$$

The transport of insulin from the liver into the blood plasma, $\mathbf{IlToIp}$ (Equation 13.2), is sent into the state **Ip** which is the total amount of insulin in plasma. The ODE of state **Ip** is kept intact from the previous model, see Equation 14 for ODE of state **Ip.**

$$\frac{d}{dt}(\mathbf{Ip}) = (\mathbf{IlToIp})\left[\frac{\mathbf{pmol/kg}}{\mathbf{min}}\right] - (\mathbf{IpToIl} + \mathbf{InsulinDegradation_{Blood}})\left[\frac{\mathbf{pmol/kg}}{\mathbf{min}}\right] \quad \text{Eq 14}$$

Insulin degradation in plasma is calculated by the function $\mathbf{InsulinDegradation_{Blood}}$, defined by product of the parameter $\mathbf{M_4}$ and the amount of insulin in the blood, state **Ip**, see Equation 14.1.

$$\mathbf{InsulinDegradation_{Blood}}\left[\frac{\mathbf{pmol/kg}}{\mathbf{min}}\right] = \mathbf{m_4}\left[\frac{\mathbf{1}}{\mathbf{min}}\right] * \mathbf{Ip}\left[\frac{\mathbf{pmol}}{\mathbf{kg}}\right] \quad \text{Eq 14.1}$$

The final part of the model consists of model variables.

### Model variables

The glycogen concentration in the liver is often measured in the unit [mmol/L]. However, meals are often defined by the unit mg per kg bodyweight, and the previous model used the same unit for e.g. glycogen. To minimize unnecessary changes from the previous model and to keep most of the parameters with similar starting guesses, we have used the same unit here in our model as well. In future model development, one should consider to change the unit to mol or kg, without the bodyweight scaling. In the model, the variable **GlycogenLiver** converts the unit from [mg/kg] to [mmol/L]. This is done through using the model input **BW**, which is the total body weight, the molecular weight of glycogen, $\mathbf{MolecularWeight_{Glycogen}}$, and the blood volume of the liver, $\mathbf{VolumeBlood_{Liver}}$, see Equation 15.1.

$$\mathbf{Glycogen_{Liver}}\ \left[\frac{\mathbf{mmol}}{\mathbf{L}}\right] = \frac{\mathbf{Gly_L}\left[\frac{\mathbf{mg}}{\mathbf{kg}}\right] * \mathbf{BW[kg]}}{\mathbf{VolumeBlood_{Liver}}\ [\mathbf{L}] * \mathbf{MolecularWeight_{Glycogen}}\left[\frac{\mathbf{g}}{\mathbf{mol}}\right]} \qquad \text{Eq 15.1}$$

$\mathbf{VolumeBlood_{Liver}}$ is a function estimating blood volume in the liver. This estimation assumes that the amount of blood in the liver is 13% of the total volume of blood in the body, calculated by function **VolumeBloodPeripheral**. There are uncertainties both in the function estimating total blood volume, **VolumeBloodPeripheral**, in data, and the assumption that the volume in the blood is 13% of the total blood volume. In several studies, the genders in populations are not homogenous or body weight and height are not documented, or both of those. To account for this uncertainty, the parameter $\mathbf{BloodLiver_{Uncertainty}}$ is introduced to calibrate volume, see Equation 15.1.1.

$$\mathbf{VolumeBloodLiver}\ [\mathbf{L}] = \quad \mathbf{VolumeBloodPeripheral[L]} * \mathbf{0.13} * \ \mathbf{BloodLiver_{uncertainty}} \qquad \text{Eq 15.1.1}$$

Model variable **VolumeBloodLiver** is linked with the total blood volume in the body estimated by the model variable **VolumeBloodPeripheral**. The function **VolumeBloodPeripheral** estimates the total amount of blood based on the model inputs **height**, **BW** and the gender of the simulated person. The function $\mathbf{VolumeBlood_{Peripheral}}$, established by Samuel B. Nadler, is a well-documented linear function estimating total blood in a human based on information of gender, length, and bodyweight. $\mathbf{Boolean_{Male}}$ and $\mathbf{Boolean_{Female}}$ is true and false statements declared in model inputs in the beginning of the simulation. The blood estimation function is kept intact from Samuel B. Nadler work [2], see Equation 15.1.2.

$$\mathbf{VolumeBlood_{Peripheral}}\ [\mathbf{L}] =$$
$$\mathbf{Boolean_{male}}\left(\mathbf{0.3669} * \left(\frac{\mathbf{height[cm]}}{\mathbf{100}}\right)^{3} + \mathbf{0.3219} * \mathbf{BW[kg]} + \mathbf{0.6041}\right) * \mathbf{BloodVolume_{uncertainty}} +$$
$$\mathbf{Boolean_{female}}\left(\mathbf{0.3561} * \left(\frac{\mathbf{height[cm]}}{\mathbf{100}}\right)^{3} + \mathbf{0.3308} * \mathbf{BW[kg]} + \mathbf{0.1833}\right) * \mathbf{BloodVolume_{uncertainty}}$$

Eq 15.1.2

The amount of insulin in the blood is often measured in the unit [pmol/L] and not the unit of the state representing insulin in blood plasma **Ip** [pmol/kg]. The unit of state **Ip** is unchanged from previous model. The model variable $\mathbf{Insulin_{Blood}}$ is used to converts the unit from [pmol/kg] to [pmol/L], see Equation 15.2.

$$\mathbf{Insulin_{Blood}}\ \left[\frac{\mathbf{pmol}}{\mathbf{L}}\right] = \frac{\mathbf{Ip}\left[\frac{\mathbf{pmol}}{\mathbf{kg}}\right] * \mathbf{BW[kg]}}{\mathbf{VolumeBloodPeripheral}\ [\mathbf{L}]} \qquad \text{Eq 15.2}$$

The amount of insulin in the blood is often measured in the unit [mg/dL] and not the unit of the state representing glucose in blood plasma **Gp** [mg/kg]. Glucose in plasma is converted from [mg/kg] to [mg/dl] with the use of model variable $\mathbf{Glucose_{Blood}}$, see Equation 15.3.

$$\mathbf{Glucose_{Blood}}\ \left[\frac{\mathbf{mg}}{\mathbf{dL}}\right] = \frac{\mathbf{Gp}\left[\frac{\mathbf{mg}}{\mathbf{kg}}\right] * \mathbf{BW[kg]}}{\mathbf{VolumeBloodPeripheral}\ [\mathbf{dL}]} \qquad \text{Eq 15.3}$$

### Data

Data from 7 clinical studies were used to train and validate the model (Table S4).

*Table S4, Summary of pre-clinical studies quantifying the metabolic flexibility of glucose homeostasis.*

| Article | Glucose metabolic flexibility status | Type of study | Population | Age [years] | BMI [kg/m$^2$] | Used for |
|---|---|---|---|---|---|---|
| Krssak *et al*. (2004) [4] | Healthy<br>T2DM | Mixed meal<br>Mixed meal | 5m/2f<br>5m/2f | 49 ± 2<br>56 ± 3 | 25.8 ± 0.9<br>26.9 ± 0.6 | Training<br>Training |
| Rothman *et al*. (1991) [5] | Healthy | Fasting | N=7 | | | Validation |
| Magnusson *et al*. (1992) [6] | Healthy<br>T2DM | Fasting<br>Fasting | 4m/1f<br>5m/2f | 61 ± 5<br>57 ± 4 | 25 ± 2<br>28 ± 1 | Training<br>Training |
| Firth *et al.* (1986) [7] | Healthy | OGTT | 1m/6f | 51±4 | 32.1 ± 2.0 | Validation |
| Lerche *et al*. (2009) [8] | Healthy | Fasting + OGTT | N=8 | 24 ± 2 | 24.4 ± 0.7 | Training |
| Taylor *et al.* (1996) [3] | Healthy | Mixed meal | 6m/2f | mean 23.6 | mean 23.1 | Validation |
| Dalla Man *et al.* (2007) [1] | Healthy | OGTT | N=204 | 56 ± 2 | 78 ± 1 kg | Training |

### Model simulations

#### Steady state

When training the model on specific flows and/or state values the optimization algorithm alters the parameter values between pre-specified reasonable bounds. In conventional mathematical ODE modelling, an initial steady-state is achieved either by starting the simulation in a perfect equilibrium, or by simulating the model to steady-state before the time period of the training occurs. To achieve a steady state the simulation first contains a standardized diet for 5 days before each study.

During these initial 5 days, meals were provided at the same times each day; breakfast 08.00, lunch 12.30 and dinner 19.00. These 3 meals per day over 5 days prior to the study were never documented in the clinical studies and their sizes were individually optimized to achieve reasonable initial values. For example, using our model we can based on glucose in plasma and/or glycogen levels estimate how much they ate.

### Model improvements

The starting point for our model development has been the Herrgårdh model [9], which in turn is an update of the Nyman model [10], which was one of the first sub-division of the glucose uptake fluxes in the original Dalla Man model [1] into specific organ fluxes. More specifically, we have taken the updated distributions between the organ fluxes from Herrgårdh *et al* 2021, but used the simplest version for the adipose tissue glucose uptake. In the full version of the Herrgårdh model, the adipose tissue glucose uptake consists of >20 ODEs, but since we are herein not considering details in the adipose tissue, we have replaced all of those with a single rate expression. This rate expression is the same as in the simplest adipose tissue sub-model in (10).

We have extended and improved the Herrgårdh model in different ways. Some of the additions that we have made to the model concerns *e.g.* the role of proteins and glycogen in glucose homeostasis at different time-scales. Another addition to the model is that we now, unlike in the previous Herrgårdh model, consider both postprandial (minutes to hours) and longer (<4 weeks) dynamic responses. These new multi-timescale capabilities of the model are all made possible by the most important addition to the model: intracellular details in the liver.

In the model, we introduce the main intracellular metabolic fluxes in the liver, which were not present in the previous models. In the original Dalla Man model, EGP was included as a separate phenomenological expression, but did not depend on the intracellular glucose concentration in the liver. In the later Herrgårdh model, the liver was introduced, in principle, but only with a single term, describing insulin-regulated glucose uptake. In our new updated liver model, we have introduced the major glucose and protein fluxes in the liver, which involve the intracellular glucose concentration, storage and breakdown of glycogen, glycolysis, and gluconeogenesis converting between glucose and pyruvate, and uptake, usage, and release of both glucose and amino acids (Fig 1D). All of these fluxes are changing both in response to a meal, and in response to more long-term changes seen for example in fasting conditions.

In the model, the main regulator of these long-term changes is glycogen, which concentration serves as a proxy for the overall energy status in the body. In other words, when glycogen concentration is high in the model, anabolic processes are upregulated and catabolic processes are downregulated, and *vice versa* when glycogen levels are low (Figure 1D, blue dashed arrow). These glycogen-dependent changes in the metabolic fluxes imply that the model can produce hypoglycaemic conditions, with plasma glucose concentration below 3.9 mM. In these hypoglycaemic conditions, the previous expression for insulin production did not work properly; therefore, that expression had to be amended.

Also on the short-term meal response level, we have improved the model, especially by adding protein dynamics (Fig 2). In the previous Herrgårdh model, only glucose and insulin were included, and any meal consisting of e.g. proteins, had to either ignore the protein content (Fig 2A,i), or convert it to glucose equivalents using a phenomenological conversion rate (Fig 2A,ii). These are two unsatisfactory options, and we have therefore added protein states which enter the gut, from which it is transported to the intestines, where proteins are digested to amino acids, which then are transported to other organs including the liver. In the other organs, amino acids are simply consumed. In the liver, amino acids are further broken down to pyruvate and other substances which enter the tricarboxylic acid (TCA) cycle, which then leads to metabolic consumption or gluconeogenesis.

### Model limitations

There are many shortcomings and assumptions in the model, many of which are due to a lack of specific clinical data that would allow and support further model development. For instance, the model does not include metabolism and crosstalk with lipids. This is a crucial shortcoming, and it means that diets such as LCFH are not simulated strictly correctly, since the ingested fat is just assumed to be consumed, with no consequence for other dynamics. This shortcoming also means that an extension of these dynamics to more long-term dynamics, involving factors such as weight change, are not possible. In model multiple intracellular metabolic reactions are highly simplified and done so through using hepatic glycogen as a homeostatic regulator. To add such more detailed intracellular models, which also incorporate in vitro experimental data, is another important future direction for the model. Furthermore, the model assumes that blood correlation between the liver and the total blood volume is constant and thus do not take into consideration of blood volume fluctuations during meal ingestion. Naturally, when doing cross-validation, training model on one population and then simulating another population, simulations of different studies give mixed results in how the model predicts reality. This is partly explained by that there are fundamental metabolic differences between populations, which highly limits the model's usability without some level of calibration data. In future work we wish to include processes describing diabetic progression and/or an appropriate mode for scaling for parameters that describes metabolic responses such as insulin resistance. Another path to solve the cross-validation issue is simply to only simulate populations where there is data on multiple meals.

In the development of our Digital twin tool no problems were observed when fitting individually to all gathered data, see Figure S2.

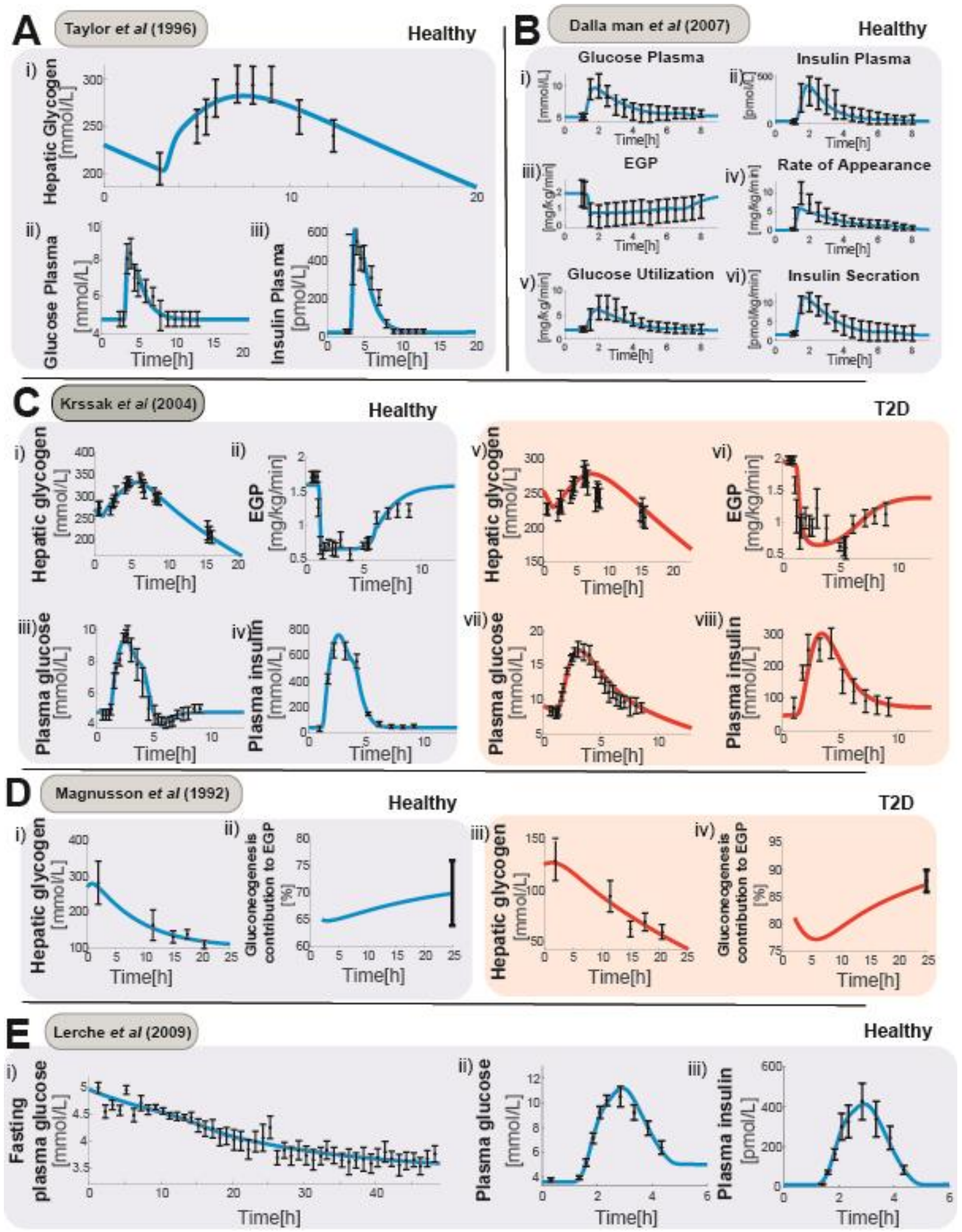

*Figure S2, Model fitted individually to several clinical studies.*

Because documented data on pure protein meals in combination with fasting are sparse, a study was conducted where participants consumed an Oral Protein Tolerance Tests (OPTT) both before and after a 48 h fasting intervention. This study utilized the model's ability in personalization and thus a smaller sample size (n=3) was chosen with a bigger focus on data resolution. The model could successfully predict glucose levels during the fast intervention and even though the model isn't trained on any pure protein meal event the model could still predict these OPTT meals and, most importantly, the differences between them. From these tests the model however predicts a bit slower glucose response from the protein drink, which underlines that in future work the protein metabolism may be advanced by following the original Dalla Man model [1] and separate all protein ingestion based on if it is consumed in a solid or liquid phase.

### References

[1] C. Dalla Man, R. A. Rizza and C. Cobelli, "Meal Simulation Model of the Glucose-Insulin System," in IEEE Transactions on Biomedical Engineering, vol. 54, no. 10, pp. 1740-1749, Oct. 2007

[2] Nadler SB, Hidalgo JH, Bloch T. Prediction of blood volume in normal human adults. Surgery. 1962 Feb;51(2):224-32. PMID: 21936146.

[3] Taylor R, Magnusson I, Rothman DL, Cline GW, Caumo A, Cobelli C, Shulman GI. Direct assessment of liver glycogen storage by 13C nuclear magnetic resonance spectroscopy and regulation of glucose homeostasis after a mixed meal in normal subjects. J Clin Invest. 1996 Jan 1;97(1):126-32. doi: 10.1172/JCI118379. PMID: 8550823; PMCID: PMC507070.

[4] Krssak M, Brehm A, Bernroider E, Anderwald C, Nowotny P, Dalla Man C, Cobelli C, Cline GW, Shulman GI, Waldhäusl W, Roden M. Alterations in postprandial hepatic glycogen metabolism in type 2 diabetes. Diabetes. 2004 Dec;53(12):3048-56. doi: 10.2337/diabetes.53.12.3048. PMID: 15561933.

[5] Rothman DL, Magnusson I, Katz LD, Shulman RG, Shulman GI. Quantitation of hepatic glycogenolysis and gluconeogenesis in fasting humans with 13C NMR. Science. 1991 Oct 25;254(5031):573-6. doi: 10.1126/science.1948033. PMID: 1948033.

[6] Magnusson I, Rothman DL, Katz LD, Shulman RG, Shulman GI. Increased rate of gluconeogenesis in type II diabetes mellitus. A 13C nuclear magnetic resonance study. J Clin Invest. 1992 Oct;90(4):1323-7. doi: 10.1172/JCI115997. PMID: 1401068; PMCID: PMC443176.

[7] Firth RG, Bell PM, Marsh HM, Hansen I, Rizza RA. Postprandial hyperglycemia in patients with noninsulin-dependent diabetes mellitus. Role of hepatic and extrahepatic tissues. J Clin Invest. 1986 May;77(5):1525-32. doi: 10.1172/JCI112467. PMID: 3517067; PMCID: PMC424555.

[8] Lerche S, Soendergaard L, Rungby J, Moeller N, Holst JJ, Schmitz OE, Brock B. No increased risk of hypoglycaemic episodes during 48 h of subcutaneous glucagon-like-peptide-1 administration in fasting healthy subjects. Clin Endocrinol (Oxf). 2009 Oct;71(4):500-6. doi: 10.1111/j.1365-2265.2008.03510.x. Epub 2008 Dec 15. PMID: 19094067.

[9] Herrgårdh T, Li H, Nyman E, Cedersund G. An Updated Organ-Based Multi-Level Model for Glucose Homeostasis: Organ Distributions, Timing, and Impact of Blood Flow. Front Physiol. 2021;12:619254-.

[10] Nyman E, Brannmark C, Palmer R, Brugard J, Nystrom FH, Stralfors P, et al. A hierarchical whole-body modeling approach elucidates the link between in Vitro insulin signaling and in Vivo glucose homeostasis. J Biol Chem. 2011;286(29):26028-41.
